# Joint effects of genes underlying a temperature specialization tradeoff in yeast

**DOI:** 10.1101/2021.03.18.436093

**Authors:** Faisal AlZaben, Julie N. Chuong, Melanie B. Abrams, Rachel B. Brem

## Abstract

A central goal of evolutionary genetics is to understand, at the molecular level, how organisms adapt to their environments. For a given trait, the answer often involves the acquisition of variants at unlinked sites across the genome. Genomic methods have achieved landmark successes in pinpointing adaptive loci. To figure out how a suite of adaptive alleles work together, and to what extent they can reconstitute the phenotype of interest, requires their transfer into an exogenous background. We studied the joint effect of adaptive, gain-of-function thermotolerance alleles at eight unlinked genes from *Saccharomyces cerevisiae*, when introduced into a thermosensitive sister species, *S. paradoxus*. Although the loci damped each other’s beneficial impact (that is, they were subject to negative epistasis), most boosted high-temperature growth alone and in combination, and none was deleterious. The complete set of eight genes was sufficient to confer ∼15% of the *S. cerevisiae* phenotype in the *S. paradoxus* background. The same loci also contributed to a heretofore unknown advantage in cold growth by *S. paradoxus*. Together, our data establish temperature resistance in yeasts as a model case of a genetically complex evolutionary tradeoff, which can be partly reconstituted from the sequential assembly of unlinked underlying loci.

**Author summary:** Organisms adapt to threats in the environment by acquiring DNA sequence variants that tweak traits to improve fitness. Experimental studies of this process have proven to be a particular challenge when they involve manipulation of a suite of genes, all on different chromosomes. We set out to understand how so many loci could work together to confer a trait. We used as a model system eight genes that govern the ability of the unicellular yeast *Saccharomyces cerevisiae* to grow at high temperature. We introduced these variant loci stepwise into a non-thermotolerant sister species, and found that the more *S. cerevisiae* alleles we added, the better the phenotype. We saw no evidence for toxic interactions between the genes as they were combined. We also used the eight-fold transgenic to dissect the biological mechanism of thermotolerance. And we discovered a tradeoff: the same alleles that boosted growth at high temperature eroded the organism’s ability to deal with cold conditions. These results serve as a case study of modular construction of a trait from nature, by assembling the genes together in one genome.

## Introduction

Understanding how organisms acquire new traits is a driving question in evolutionary biology. Many traits of interest are adaptations, meaning they provide a fitness benefit to the organism which has driven their rise to high frequency in the population. Genomic methods often find a slew of unlinked changes at the DNA level that associate with a given adaptive trait (Burke and Rose, 2009; Iranmehr et al., 2021; Pardo-Diaz et al., 2015; Sella and Barton, 2019). For any one candidate locus, gold-standard validation experiments will then swap alleles between taxa and test for an impact on phenotype, at genic (Castro et al., 2019; Chan et al., 2010; Kryazhimskiy et al., 2014; Linnen et al., 2013, 2009; Xie et al., 2019) and sub-genic (Anderson et al., 2016; Bridgham et al., 2009; Escudero et al., 2020; Finnigan et al., 2012; Lindsey et al., 2013; Liu et al., 2018; Palmer et al., 2015, 2015; Pillai et al., 2020; Toprak et al., 2011; Weinreich et al., 2006) levels of resolution. Though elegant and rigorous, this focused approach on a given gene will by necessity leave polygenic mechanisms of the trait less well characterized.

For a more complete picture of a complex adaptation, we would establish how multiple underlying genes work together, including their interdependence and their joint ability to recapitulate the phenotype. Such questions have come within reach in laboratory evolution, with particular emphasis on genomic inferences of epistasis between unlinked adaptive loci (Aggeli et al., 2020; Blount et al., 2012; Bons et al., 2020; Buskirk et al., 2017; Csilléry et al., 2018; Fisher et al., 2019; Good et al., 2017; Johnson et al., 2021; Kryazhimskiy et al., 2014; Weber et al., 1999). In a handful of cases, adaptive multi-gene interactions from a lab evolution have been verified experimentally by allelic replacement (Chou et al., 2011; Khan et al., 2011). To date, validating these principles in the context of evolution from the wild has posed a key challenge (although see Marcusson et al., 2009; Neverov et al., 2015; Roop et al., 2016).

To study complex genetic mechanisms in adaptation, we set out to use natural variation in *Saccharomyces* yeasts as a model. *S. cerevisiae* strains, from the wild and the lab, grow at temperatures up to 41°C (Gonçalves et al., 2011; Salvadó et al., 2011; Sweeney et al., 2004). All other species in the *Saccharomyces* clade, which diverged from a common ancestor ∼20 million years ago, grow poorly at high temperatures, though many outperform *S. cerevisiae* in the cold (Hittinger, 2013). In previous work (Weiss et al., 2018) we developed a genomic version of the reciprocal hemizygosity test to dissect thermotolerance, using *S. paradoxus*, the closest sister species to *S. cerevisiae*, as a representative of the inferred ancestral state. Derived alleles of the mapped genes in *S. cerevisiae*, when tested individually for their marginal effects, were partially necessary or sufficient for thermotolerance, or both (Weiss et al., 2018), and their sequences exhibit evidence for positive selection in *S. cerevisiae* (Abrams et al., 2021; Weiss et al., 2018). But how these genes work together has remained unknown. We thus aimed to investigate the extent to which unlinked thermotolerance loci assembled in the same background would explain the trait, and whether and how these genes would depend on one another for their effects. We expected that any answers could also help elucidate other facets of the mechanism and the evolutionary history of thermotolerance.

## Results

### Combining thermotolerance loci to reconstitute a complex trait

We previously mapped eight genes with pro-thermotolerance alleles in *S. cerevisiae* (Table 1; Weiss et al., 2018). To explore the joint function of these unlinked loci, we introduced the alleles of all eight from DBVPG1373, a soil isolate of *S. cerevisiae* from the Netherlands, into Z1, a strain of the sister species *S. paradoxus*, isolated from an English oak tree (Liti et al., 2009). Our approach used a stepwise set of gene replacements. With CRISPR/Cas9 we introduced the *S. cerevisiae* allele of the promoter and coding region of a given gene into the endogenous location in wild-type *S. paradoxus*; we used the resulting strain as a background for the replacement of the *S. paradoxus* allele of the next gene by that of *S. cerevisiae*; and so on until all eight genes were swapped into one genome (Figure 1, bottom).

**Table 1.** Genes contributing to thermotolerance divergence between *Saccharomyces cerevisiae* and *S. paradoxus*. GO, Gene Ontology biological process. The last column reports amino acid divergence between *S. cerevisiae* DBVPG1373 and *S. paradoxus* Z1.

| Gene | GO terms | Location | Description | Protein Length (residues) | Protein Identity (%) |
| --- | --- | --- | --- | --- | --- |
| *AFG2/YLR397C* | ribosomal large subunit biogenesis | Ch. XII<br>912550 -<br>914892 | Essential for pre-60S maturation and release of several pre-ribosome maturation factors | 780 | 93.6 |
| *APC1/YNL172W* | mitotic cell cycle | Ch. XIV<br>310636 -<br>315882 | Largest subunit of the Anaphase-Promoting Complex; a ubiquitin-protein ligase required for degradation of anaphase inhibitors | 1748 | 90.6 |
| *CEP3/YMR168C* | mitotic cell cycle | Ch. XIII<br>597332 -<br>599158 | Essential kinetochore protein; component of the CBF3 complex that binds the CDEIII region of the centromere | 575 | 93.2 |
| *DYN1/YKR054C* | mitotic cell cycle | Ch. XI<br>535647 -<br>547925 | Heavy chain dynein; microtubule motor protein; required for anaphase spindle elongation | 4092 | 87.5 |
| *ESP1/YGR098C* | mitotic cell cycle | Ch. VII<br>682566 -<br>687458 | Separase/separin; aids in the dislocation of cohesin from chromatin and sister chromatin segregation. | 1630 | 89.9 |
| *MYO1/YHR023W* | mitotic cell cycle | Ch. VIII<br>151666 -<br>157452 | Type II myosin heavy chain; required for cytokinesis and cell separation | 1928 | 91.9 |
| *SCC2/YDR180W* | mitotic cell cycle | Ch. IV<br>821295 -<br>825776 | A complex required for loading of cohesin complexes onto chromosomes; involved in establishing sister chromatid cohesion during DSB repair via histone H2AX; subunit of cohesin loading factor (SCC2p-Scc4p) | 1493 | 88.2 |
| TAF2/YCR042C | chromatin<br>binding | Ch. III 201174<br>- 205397 | TFIID subunit; involved in RNA polymerase II<br>transcription initiation | 1407 | 89.8 |

**Figure 1.**
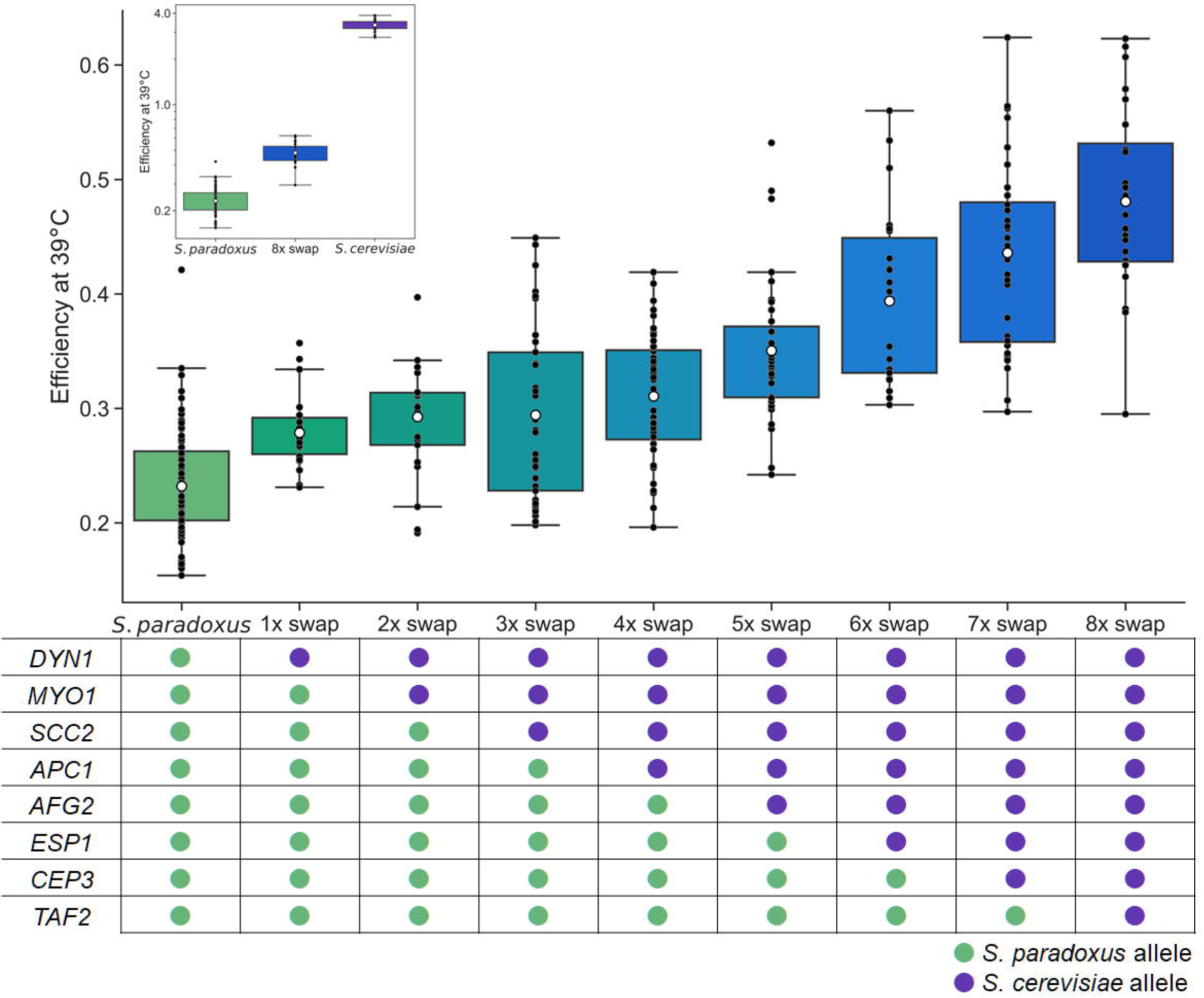
*S. cerevisiae* alleles of thermotolerance loci jointly improve growth at high temperature. In each plot, the *y*-axis reports growth efficiency at 39°C, the cell density after a 24-hour incubation as a difference from the starting density. In the main plot, each column reports data from a transgenic *S. paradoxus* strain harboring *S. cerevisiae* alleles of the indicated thermotolerance loci, or the wild-type *S. paradoxus* progenitor. At bottom, each cell reports the genotype at the indicated locus in the indicated strain. The inset shows purebred wild-types and the *S. paradoxus* strain harboring all eight thermotolerance loci from *S. cerevisiae* (8x swap); the *y*-axis is log-scaled. Black points report individual biological replicates and white dots report means. Boxes span the interquartile range. Whiskers are 1.5 times the interquartile range. Statistical analyses are reported in Table S1A.

We first assayed growth of the eight-gene transgenic and the wild-type parental species at 39°C, with biomass accumulation (growth efficiency; the increase in optical density after 24 hours relative to that at the start) as a readout of strain performance. The results revealed an advantage of 2.07-fold attributable to the eight *S. cerevisiae* alleles in the *S. paradoxus* background (Figure 1, inset and Table S1A), and no such effect in 28°C control conditions (Figure S1). This joint phenotype recapitulated 15% of the trait divergence between wild-type *S. cerevisiae* and *S. paradoxus* at 39°C (Figure 1, inset), with comparable results at temperatures down to 37°C (Figure S2). These data make clear that the full genetic architecture of *S. cerevisiae* thermotolerance must involve more loci than the eight we have manipulated here—highlighting the potential for high genetic complexity of this trait divergence between species.

### Negative epistasis among thermotolerance genes

We hypothesized that *S. cerevisiae* thermotolerance determinants might depend on one another to confer their effects. We sought to test this at the whole-gene level, treating the allele of each thermotolerance locus (including the promoter and coding region) as a module, and investigating the interactions between them. The *S. cerevisiae* allele of each module, when introduced on its own into *S. paradoxus*, was sufficient for a <1.4-fold benefit in biomass accumulation at 39°C, as expected (Figure S3 and Weiss et al., 2018). We summed these measurements to yield an expected phenotype under the assumption of independent gene function, which we compared to the true measurement from the *S. paradoxus* strain harboring all eight thermotolerance loci from *S. cerevisiae*. The latter came in significantly below the estimate from the model assuming independence (Figure 2A). Thus, the combination of all eight genes was subject to negative epistasis, improving growth at high temperature to an extent less than the sum of its parts.

**Figure 2.**
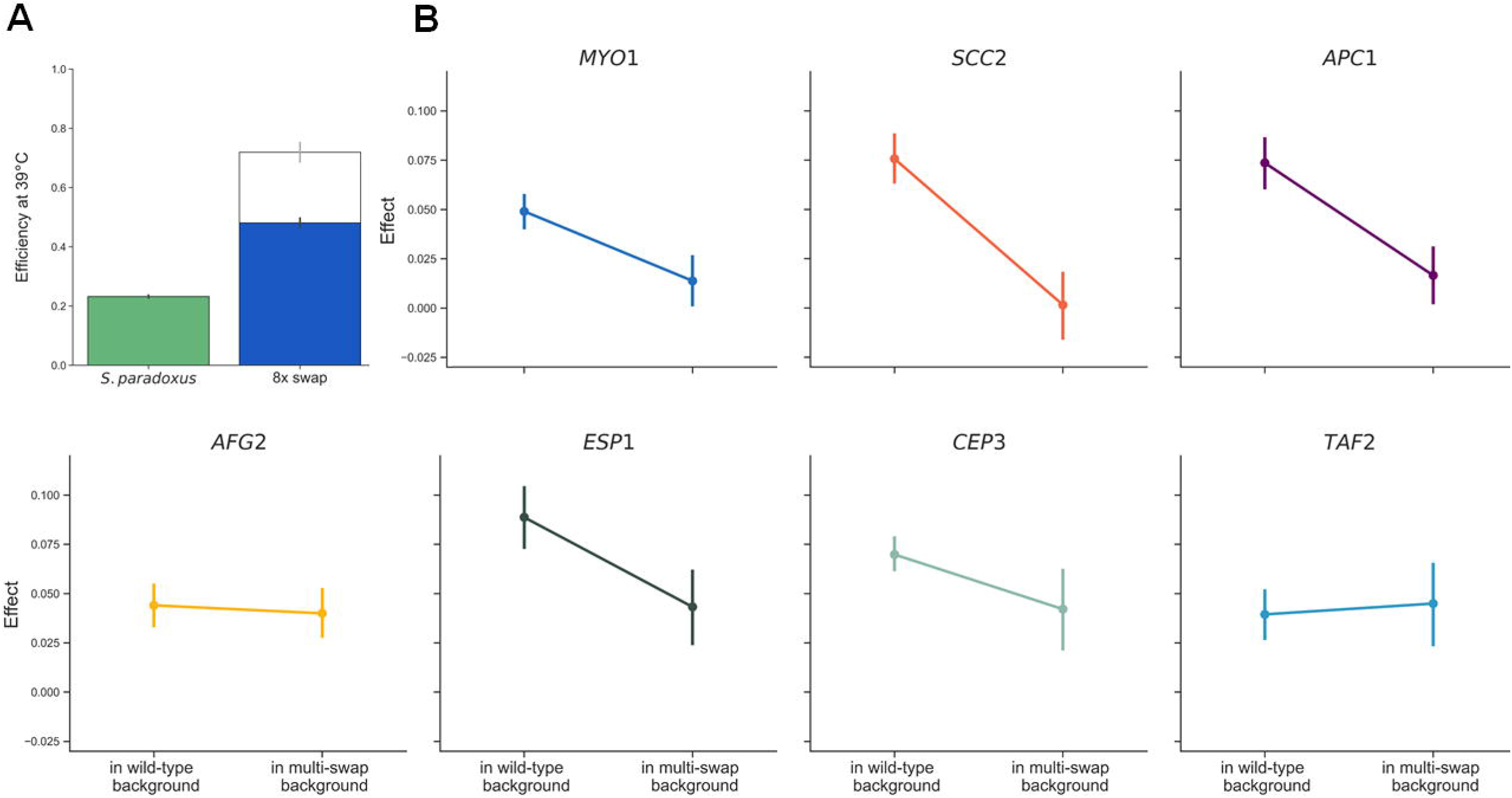
Negative epistasis among *S. cerevisiae* alleles of thermotolerance loci. (A) Solid bars report growth efficiency at 39°C for purebred *S. paradoxus* and the *S. paradoxus* strain harboring all eight thermotolerance loci from *S. cerevisiae* (8x swap). In the right column, the hollow extension reports the sum of the efficiencies at 39°C of *S. paradoxus* strains harboring individual thermotolerance loci from *S. cerevisiae*, from Figure S3. (B) In a given panel, the right-hand point shows the effect, on efficiency at 39°C, of the *S. cerevisiae* allele of the indicated gene when introduced into a transgenic also harboring *S. cerevisiae* alleles of other genes, in the series of **Figure 1**. The left-hand point shows the analogous quantity when wild-type *S. paradoxus* was the background. Error bars report 68% confidence intervals.

We next turned to the strains we had made in the service of the eight-gene transgenic. These harbored *S. cerevisiae* modules of one thermotolerance gene in the *S. paradoxus* background, two genes, three genes, and so on (Figure 1, bottom). We considered this strain panel as an arbitrary trajectory through a gene-wise genetic landscape from the *S. paradoxus* wild-type to the eight-locus swap. Though it represented just one of tens of thousands of possible paths, we anticipated that it could help inform our understanding of the genetic architecture of thermotolerance.

In assays of biomass accumulation at 39°C in each strain of the panel, we saw a generally monotonic relationship between thermotolerance and the number of *S. cerevisiae* gene modules introduced into *S. paradoxus* (Figure 1 and Table S1A). No such pattern emerged from a 28°C control (Figure S1 and Table S1B). For a quantitative analysis, we converted the phenotypic profiles in the strain set to effect sizes, each reporting the impact of the *S. cerevisiae* allele of a gene on thermotolerance, in the chimeric background into which it was introduced (Figure 2B). Inspection of this metric confirmed that each *S. cerevisiae* gene addition boosted the phenotype along the path to the eight-fold transgenic, in most cases significantly so (Table S1A), with one exception. The pro-thermotolerance function of *S. cerevisiae SCC2*, discernable when it was introduced on its own into *S. paradoxus* (Figure S3), was either below our detection limit or absent altogether in combination with *S. cerevisiae DYN1* and *MYO1* (Figures 1 and 2B). Despite this potential case of masking epistasis, the remaining trend for beneficial effects by *S. cerevisiae* alleles as we built up the eight-locus strain was highly non-random (binomial *p* = 0.03). In no case did we observe a defect from swapping in an *S. cerevisiae* allele, meaning we had no evidence for sign epistasis in this system.

We also used our multi-genic strain panel, alongside single-gene transgenics in *S. paradoxus*, to compare *S. cerevisiae* allele-replacement effects across backgrounds. In most cases, a given thermotolerance gene from *S. cerevisiae* had less impact in the presence of other *S. cerevisiae* loci than when tested on its own in *S. paradoxus* (Figure 2B). This finding confirmed the negative (magnitude) epistasis between the *S. cerevisiae* alleles of thermotolerance genes that we had inferred with the eight-gene transgenic (Figure 2A). We noted that the damping of allelic effect by genetic background was most apparent at thermotolerance genes annotated in chromosome segregation/mitosis (*DYN1, MYO1, SCC2, APC1*, and *ESP1;* Figure 2B). By contrast, introducing the *S. cerevisiae* allele of *TAF2* or *AFG2*, involved in transcription and translation respectively, drove nearly the same benefit on its own in *S. paradoxus* and in the respective multi-swap chimera (Figure 2B).

Together, these genetic data characterize an example path toward thermotolerance of incremental advances from *S. cerevisiae* gene modules—most limiting each other’s effects to some extent, but without frank deleterious consequences.

### Thermotolerance loci improve viability only during active growth

We next aimed to investigate cellular mechanisms of thermotolerance, using as a tool the strain with all eight of our focal genes from *S. cerevisiae* replaced into *S. paradoxus*. We focused on cell viability, as assayed by counts of colony-forming units (CFUs) from aliquots of liquid culture at 39°C. In a first characterization of strain performance in growing cultures under this setup, *S. paradoxus* cells were much less viable than those of *S. cerevisiae*, across a range of warm temperatures (Figure S4), as expected. We anticipated that *S. cerevisiae* alleles of our thermotolerance loci would rescue this phenotype, at least in part. This prediction bore out in our CFU assays from actively growing cultures: at 39°C, the eight-gene transgenic survived 7-fold better than did its *S. paradoxus* progenitor (Figure 3A). An analogous test of wild-type *S. cerevisiae* revealed three logs higher viability than that of *S. paradoxus* during active growth at 39°C (Figure 3A).

**Figure 3.**
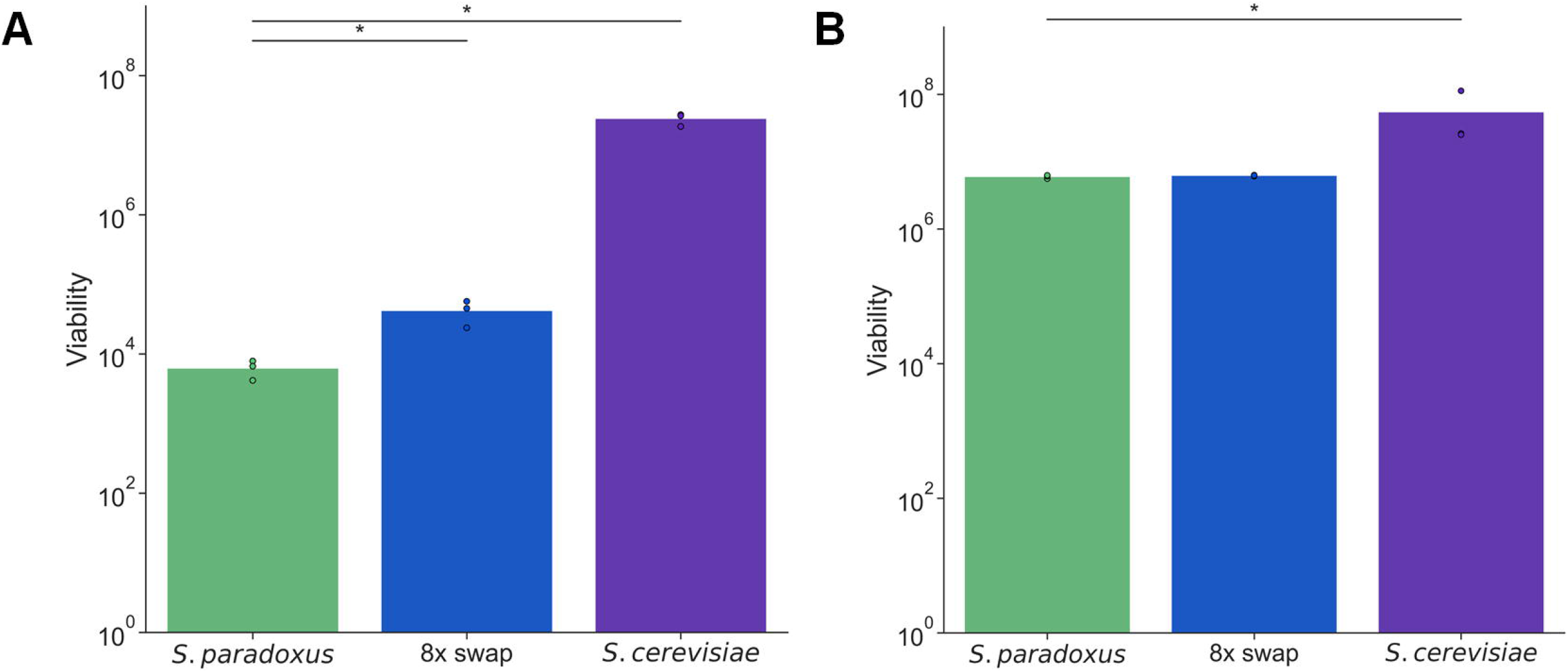
*S. cerevisiae* alleles of thermotolerance loci jointly improve heat survival during growth but not in stationary phase. In a given panel, each column reports viability after heat treatment of the wild-type of the indicated species, or the *S. paradoxus* strain harboring eight thermotolerance loci from *S. cerevisiae* (8x swap). The *y*-axis reports the number of colonies formed on solid medium from 1 mL of heat-treated liquid culture in (A) logarithmic growth or (B) stationary phase, normalized by turbidity. Points and bar heights report individual biological replicates and their means, respectively. *, Wilcoxon *p* ≤ 0.05.

The poor performance of strains of *S. paradoxus* origin in the CFU assay (Figure 3A) was much bigger in magnitude than that seen in our measurements of biomass accumulation (Figure 1), in growing cultures. Such a discrepancy suggests that some cells with *S. paradoxus* genotypes manage to divide early in the heat treatment and then ultimately die, contributing at the end of the incubation to measured biomass but not viability. Qualitatively, however, both experiments led to the same conclusion: *S. cerevisiae* alleles of thermotolerance loci recapitulate part but not all of the advantage by *S. cerevisiae* relative to *S. paradoxus* during active growth at 39°C. We detected no viability differences between strains during active growth at 28°C (Figure S5A).

To gain insight into why *S. paradoxus* cells die at high temperature, we took account of the role in mitosis for most of our thermotolerance genes (Table 1). We hypothesized that failure of the *S. paradoxus* cell growth machinery was the proximal cause of death for this species at 39°C. If so, we expected that the underlying alleles would not be a liability if cells did not enter the cell cycle in the first place. As a test of this notion, we retooled our viability assay to start by incubating a liquid culture at a permissive temperature until it reached stationary phase (when nutrients are exhausted and cell division arrests). We then switched these non-growing cultures to 39°C, and finally took aliquots for assays of CFUs. The results revealed that, when exposed to heat as a stationary-phase culture, *S. paradoxus* survived nearly as well as did *S. cerevisiae* (Figure 3B), in contrast to the many logs of difference between the species during active growth (Figure 3A). Likewise, transgenesis of our eight thermotolerance genes had no impact on high temperature survival in stationary phase (Figure 3B). Viability experiments on stationary-phase cultures at 28°C also found no difference between strains (Figure S5B).

These viability data show that the thermotolerance defect of *S. paradoxus* alleles from our eight-gene transgenic strain only manifests in actively growing cells, consistent with the hallmarks of cell cycle breakdown seen in microscopy assays of *S. paradoxus* at 39°C (Weiss et al., 2018). Together, our results support a model in which passage through mitosis itself is lethal at high temperature for *S. paradoxus*, whereas cells in an arrested state are protected from damage and death. *S. cerevisiae*, meanwhile, grows and divides successfully at 39°C, owing in part to its thermotolerant mitotic genes.

### An evolutionary tradeoff in tolerance of extreme temperatures

Given that *S. cerevisiae* is unique within its clade for its ability to grow at high temperatures, we anticipated that this trait could have evolved as part of a tradeoff, and that cold tolerance would be a logical potential opposing character. Consistent with this picture, we observed generally better cold resistance across a panel of wild *S. paradoxus* relative to environmental isolates of *S. cerevisiae* (Figure S6). No such difference is detectable at 28°C (Weiss et al., 2018). We reasoned that thermotolerance alleles at our eight focal genes could contribute to the poor growth by *S. cerevisiae* in the cold. Indeed, our eight-fold transgenic strain grew significantly worse than did wild-type *S. paradoxus* at 4°C (Figure 4). With respect to biomass accumulation, this strain recapitulated 15% of the divergence between the wild-type species at 4°C—paralleling the analogous quantity at 39°C (Figure 1), and establishing antagonistic pleiotropy by *S. cerevisiae* alleles at our focal genes.

**Figure 4.**
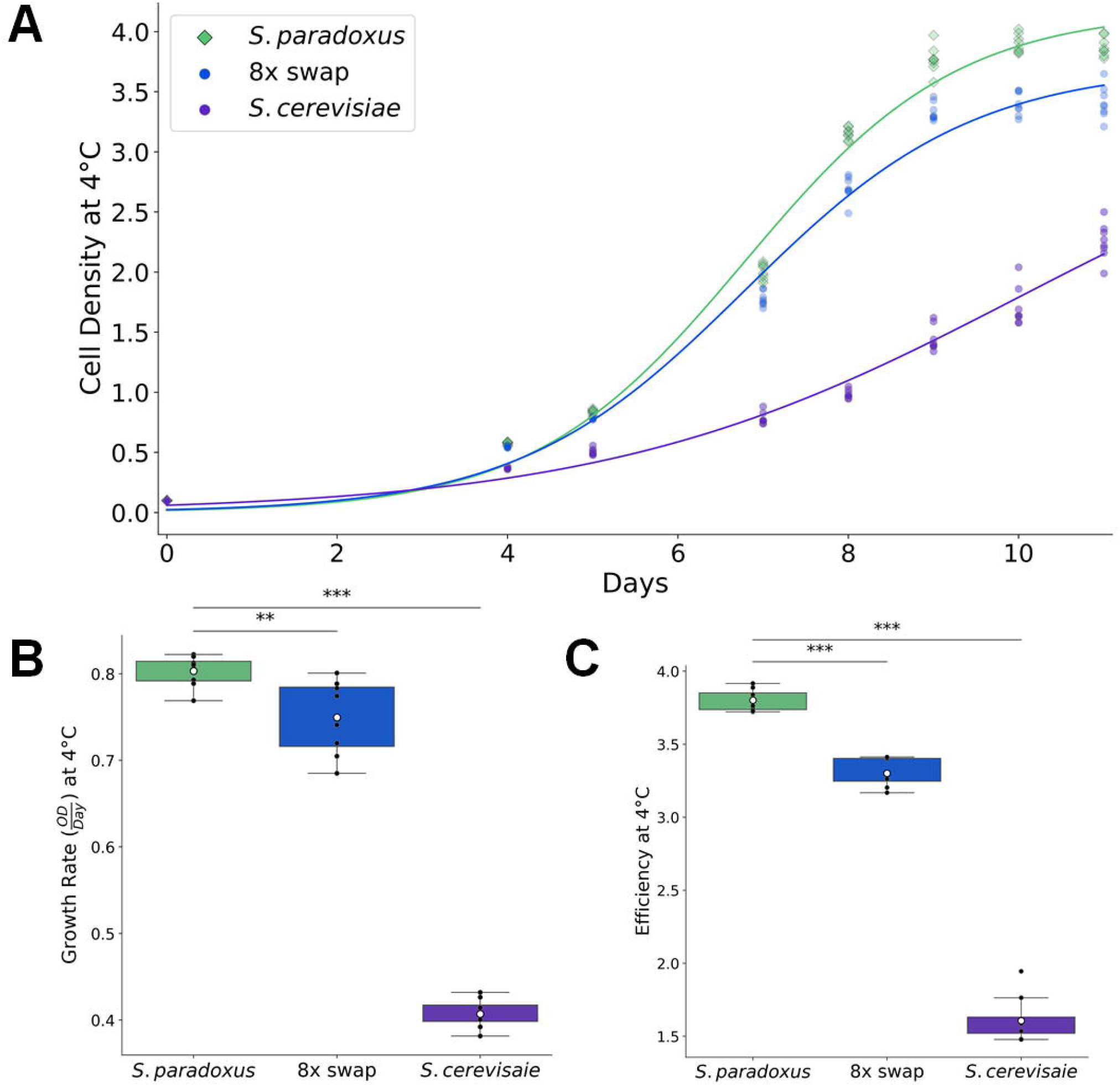
*S. cerevisiae* alleles of thermotolerance loci jointly compromise growth in the cold. (A) Each trace reports a timecourse of growth at 4°C of the wild-type of the indicated species, or the *S. paradoxus* strain harboring eight thermotolerance loci from *S. cerevisiae* (8x swap). For a given strain, points on a given day report biological replicates; lines report the average fit from a logistic regression across replicates (Table S2). (B) The *y*-axis reports growth rate, in units of cell density (optical density, OD) per day, from the average logistic fit of the timecourse in (A) for the indicated strain. (C) The *y*-axis reports, for day 10 of the timecourse in 1.for the indicated strain, growth efficiency, the cell density after a 10-day incubation at 4°C as a difference from the starting density. In (B) and (C), points report individual biological replicates. Boxes show the median and span the interquartile range of the data (IQR, 25%-75%); whiskers are 1.5 times the interquartile range, and do not report outliers. ** and ***, Wilcoxon *p* ≤ 0.004 and *p* ≤ 0.0005, respectively.

We hypothesized that thermotolerance loci could contribute to growth in the cold by mechanisms that would differ from the genetics we had elucidated at high temperature. In analysis of individual genes, only the *S. cerevisiae* allele of *DYN1* reduced biomass accumulation markedly at 4°C when introduced into *S. paradoxus* on its own, though a number of other genes had smaller effects (Figure S7). Likewise, across our panel of combinatorial gene replacements, most strains roughly phenocopied the single-gene *DYN1* swap in terms of growth efficiency (Figure S8). This contrasted with the stairstep-like trend of increasing thermotolerance as *S. cerevisiae* alleles were stacked in the *S. paradoxus* background (Figure 1). We thus considered *DYN1* as a central player in the cold sensitivity of *S. cerevisiae*, against a backdrop of more subtle contributions from other loci, alone and in combination.

In viability assays during growth at 39°C, we had found robust divergence in survival between *S. cerevisiae* and *S. paradoxus*, as well as survival effects of variation at our thermotolerance loci (Figure 3A). It remained an open question whether the genetics of cold tolerance would have effects on cell death. In viability assays we saw very little difference between *S. cerevisiae, S. paradoxus*, and our eight-gene transgenic in the *S. paradoxus* background, in terms of CFUs after incubation at 4°C (Figure S9). This result underscores the difference in mechanism between cryotolerance and thermotolerance in this system, and establishes that *S. cerevisiae* alleles (at thermotolerance loci and others) slow growth in the cold rather than killing cells outright.

## Discussion

Evolution often uses variants at genes scattered throughout the genome to build an adaptive trait. The complete set of these loci, once in hand, can reconstitute the trait in an exogenous background, which is the ultimate goal for many biomedically and industrially relevant characters. Along the way, the underlying genes can shed light on the process by which a trait arose, which may involve events from millions of years ago. In this work, we have used thermotolerance, a model fitness-relevant character whose underlying genes have been under positive selection in environmental isolates of *S. cerevisiae* (Abrams et al., 2021; Weiss et al., 2018), to explore genetic, biological, and evolutionary mechanisms of adaptation.

Introducing eight unlinked genes from *S. cerevisiae* into *S. paradoxus*, we reconstituted ∼15% of the difference in thermotolerance between the respective purebred species. Much of the architecture for this trait thus remains unmapped, likely due in part to limitations of coverage and power in our original reciprocal hemizygosity scan (Weiss et al., 2018), and to its restriction to the nuclear genome. Incisive experiments have established the role of mitotype in temperature tolerance as it differs between *S. cerevisiae* and either *S. paradoxus* or a farther-diverged species of the complex, *S. uvarum* (Baker et al., 2019; Hewitt et al., 2020; Li et al., 2019; Špírek et al., 2014). Thus, mitochondrial variants presumably make up much of the heritability missing from our current gene set. And *S. cerevisiae* alleles of the nuclear-encoded genes that we study here could well depend on the *S. cerevisiae* mitochondrial genome for their full effect.

In our focus on nuclear-encoded thermotolerance factors, we found that most *S. cerevisiae* alleles improved high-temperature growth, when tested on their own in *S. paradoxus* or in a chimeric multi-swap background. In principle, these effects could be limited by partial incompatibilities in *S. paradoxus*—*i*.*e*. a given gene could have made an even bigger contribution to thermotolerance in the presence of other functional partners from *S. cerevisiae*. Even in such a case, the qualitative conclusion from our own data would remain: *S. cerevisiae* alleles mostly help and do not hurt at high temperature, alone or combined. This profile provides an intriguing contrast to the sign epistasis often detected between adaptive amino acid changes within a given protein, as they boost fitness in some combinations and exert a frank defect in others (Starr and Thornton, 2016; Weinreich et al., 2005). Since we have not surveyed all possible subsets of our mapped thermotolerance genes, and more remain unidentified, we cannot rule out the possibility of toxic interactions at high temperature between some *S. cerevisiae* versions of unlinked loci when they come together. That said, our thermotolerance data as they stand conform to the idea that combining adaptive variants across unlinked sites will be less constrained than, say, repacking a given protein (Toprak et al., 2011). This model is especially compelling in the search to understand evolutionary dynamics in sexual systems. If a gene module can recombine into most backgrounds and confer a benefit, it would speed the advance toward fitness of the population as a whole.

We found that the magnitude of the phenotypic effect of our focal genes depended on genetic background. For our analyses of epistasis, we focused on thermotolerance, under the assumption that this trait or its correlates have mattered for fitness in wild *S. cerevisiae*, and thus that genetic interactions could have been at play during the adaptation process. We found ample evidence for negative epistasis, with the *S. cerevisiae* alleles of some of our unlinked loci obscuring the thermotolerance effects of others, in replacements into *S. paradoxus*. Negative epistasis has been a linchpin of classical genetic studies placing unlinked loss-of-function mutants in pathways (Avery and Wasserman, 1992; Roth et al., 2009), and their genomic equivalent, reverse-genetic double-mutant screens (Baryshnikova et al., 2013; Shen et al., 2017; Tutuncuoglu and Krogan, 2019; Wong et al., 2016). Negative epistasis also features in the polygenic architectures of putatively neutral and disease traits that vary in populations (Gusareva and Van Steen, 2014; Mackay, 2014; Ritchie, 2015). Our thermotolerance system complements these scenarios, in that we focus on alleles that act as gains of function, at least at high temperature (Weiss et al., 2018). Under the latter condition, we detected the most marked negative epistasis among thermotolerance genes with annotations in chromosome segregation and cell division. We can speculate that the *S. cerevisiae* allele at any such locus, as it rescues one aspect of cell division at 39°C, also pulls up the function of other parts of the mitotic machinery to some extent, *e*.*g*. by stabilizing heteromeric protein complexes. *S. cerevisiae* alleles of additional genes, introduced into such a background, would make less dramatic improvements to the phenotype than would be expected from their respective single-gene allele-swap strains.

Importantly, a sizeable literature has described negative epistasis between unlinked adaptive loci in laboratory evolution experiments, especially in microbes (Aggeli et al., 2020; Blount et al., 2012; Bons et al., 2020; Buskirk et al., 2017; Csilléry et al., 2018; Fisher et al., 2019; Good et al., 2017; Johnson et al., 2021; Kryazhimskiy et al., 2014; Tenaillon et al., 2012; Weber et al., 1999), where suites of unlinked variants have been validated in terms of effects on fitness (Chou et al., 2011; Khan et al., 2011). A chief emphasis here has been on evolutionary dynamics, in that negative epistatic interactions limit the availability of mutational steps with big fitness effects that would otherwise speed adaptation (Starr and Thornton, 2016; Weinreich et al., 2005). This principle plays out in part through diminishing-returns epistasis (Kryazhimskiy et al., 2014; Wiser et al., 2013), the difficulty of improving the fitness of somewhat-fit backgrounds, which can be explained by heightened susceptibility of fast-growing strains to pleiotropic effects of new mutations (Johnson et al., 2019). Though the latter has been a landmark of the recent literature, we do not consider it an immediately relevant model for our own data because of our focus on backgrounds that perform very poorly at 39°C. In more advanced stages of an allele-swap progression, which would reconstitute more of the yeast thermotolerance phenotype, we would expect genes to interact under the diminishing-returns mechanism.

Along with our insights into the genetics of thermotolerance, we discovered a cold resistance defect in *S. cerevisiae* relative to *S. paradoxus*, complementing previous temperature profiling using other assays across the genus (Hittinger, 2013; Salvadó et al., 2011). And we were able to pinpoint the genetic basis of this phenotype, at least in part, to pro-thermotolerance *S. cerevisiae* alleles at our focal genes. The latter could be attributed largely to *S. cerevisiae DYN1*, which elicited the most dramatic drop in cold tolerance when tested on its own in *S. paradoxus*, and overshadowed the effects of other loci in multi-gene swap backgrounds. This echoes the known breakdown of *S. cerevisiae DYN1* protein function below 8°C, as distinguished from the mammalian ortholog (Hong et al., 2016). The emerging picture is that *S. cerevisiae* harbors alleles of *DYN1* (and other genes) that have been adaptive in its ancestral niche and yet act as a liability in the cold; other deleterious effects could also be at play, in conditions beside those studied here. Given that *S. cerevisiae* also tolerates ethanol better than the rest of its clade (Herbst et al., 2017; Williams et al., 2015), one appealing model posits that its unique ability to ferment drove *S. cerevisiae*’s origin as a species (Goddard, 2008), generating heat and ethanol at levels that kill off its microbial competitors. If so, our data strongly suggest that *S. cerevisiae* could not achieve this adaptation as a temperature generalist, and had to make a tradeoff by acquiring alleles that eroded the capacity to deal with cold.

## Acknowledgements

The authors thank Dmitri Petrov for motivating the analysis of the thermotolerance of stationary-phase cultures; Abel Duarte and Jeff Skerker for direction and advice with molecular biology; Carly Weiss and Jeremy Roop for other helpful discussions; and David Savage for his generosity with lab facilities and resources.

## Methods

### Strain construction

Strains used in this work are listed in Table S3. To study combinations of *S. cerevisiae* alleles in an *S. paradoxus* background, we used as the latter the wild-type homozygous diploid *S. paradoxus* Z1, originally isolated from tree bark in the UK (Liti et al., 2009). We used our CRISPR/Cas9 method (Weiss et al., 2018), essentially as described but with slight modifications detailed below, to replace both copies of a given thermotolerance locus, at the endogenous location, with the allele from *S. cerevisiae* DBVPG1373, a soil isolate from the Netherlands (Liti et al., 2009). To build the eight-gene transgenic, we started with the single-gene transgenic in the Z1 background harboring the *S. cerevisiae* allele of *DYN1* from Weiss et al., 2018. We introduced *S. cerevisiae* alleles at additional genes in an iterative series of transformations; after each, we cultured an isogenic stock from a single colony for Sanger sequence confirmation and storage, and, where appropriate, we used this stock as input into the next transformation.

A given transformation involved donor DNA and constructs encoding guide RNAs for Cas9 to target replacement of the *S. paradoxus* allele with that from *S. cerevisiae* at either one or two loci, which Sanger sequencing then verified to be successful at one or both. For each locus we used designs of guide RNAs from Weiss et al., 2018 that targeted, for double-strand breaks, the endogenous Z1 allele by Cas9; one guide targeted a site ∼1000bp upstream of the coding start of the gene of interest (or the 3’ end of the closest upstream gene) in the Z1 genome and the other guide targeted a site near the coding stop. Cas9 editing proceeded as described (Weiss et al., 2018). Briefly, for each transformation step of strain construction, one or two guide RNA pairs, targeting one or two loci respectively, were cloned into plasmid pBC712, which also encodes *Streptococcus pyogenes* Cas9. Next, DNA to serve as a repair template was generated for each relevant locus, via PCR from *S. cerevisiae* DBVPG1373, with 90bp primers that contained 70bp of sequence homologous to *S. paradoxus* Z1 on each side of the amplified DNA product. Finally, plasmid and repair template were transformed into the Z1 descendant as described (Weiss et al., 2018) at a ratio of 0.3 to 3.5 based on the length of the donor DNA, equivalent to 10^12^ dsDNA molecules for 10 μg of plasmid. Putative transformants were purified and sequence-verified.

### Growth assays

Measurements of biomass accumulation (growth efficiency) at 39°C in Figure 1 were done essentially as described (Weiss et al., 2018) with modifications as follows. For a given day’s worth of experiments, wild-type *S. paradoxus* Z1 and one or more other strains of interest were streaked onto a yeast peptone dextrose (YPD) plate from a -80°C freezer stock and incubated at 28°C for 2 days. 2-8 colonies of a given strain were each inoculated separately in 5mL of liquid YPD and grown for 24 hours at 28°C with shaking at 200rpm to saturation; we refer to the cultures at this stage as pre-cultures. Each such replicate pre-culture was back-diluted into 10mL of YPD to achieve an OD_600_ of 0.05; incubated at 28°C until it reached an OD_600_ of 0.4-0.8; back-diluted again to OD_600_ of to achieve an OD_600_ of 0.1; and then incubated for 24 hours at 39°C. We tabulated the difference in OD_600_ between the final and initial timepoints across this 24-hour incubation for each culture. This procedure, from streaking on solid medium through inoculation, heat treatment, and biomass measurement, was repeated at least four times for each strain in the analysis of Figure 1. The resulting vector of biomass measurements across all replicates from all days for each strain was compared to that for each other strain with a one-tailed Wilcoxon test in Table S1. Multiple testing was corrected for using the Benjamini-Hochberg method.

Measurements of biomass accumulation at 28°C in Figure S1 were done as described (Weiss et al., 2018). Strains were streaked on solid plates and one colony per strain was pre-cultured in liquid at 28°C as above. Each such saturated pre-culture was back-diluted to achieve an OD_600_ of 0.05 and grown for an additional 5.5 hours at 28°C until it reached logarithmic phase. We transferred cells from each such pre-culture, and YPD, to five replicate wells of a 96-well plate, with volumes sufficient to yield a total volume of 150 μL per well at an OD_600_ of 0.02. The plate was covered with a gas-permeable membrane (Sigma) and incubated with orbital shaking in an M200 plate reader (Tecan, Inc.) at 28°C for 24 hours. We tabulated the difference in OD_600_ between the final and initial timepoints across this 24-hour incubation for each replicate culture. The vector of these replicate measurements for each strain was compared to that from *S. paradoxus* with a two-tailed Wilcoxon test. Multiple testing was corrected for using the Benjamini-Hochberg method.

For measurements of biomass accumulation at 37°C -39°C in Figure S2, strains were streaked, and three colonies of a given strain were pre-cultured in liquid as above. Each such replicate liquid culture was back-diluted into 10mL of YPD to achieve an OD_600_ of 0.05; incubated at 28°C until it reached an OD_600_ of 0.4-0.8; back-diluted again to OD_600_ of to achieve an OD_600_ of 0.1; and then incubated for 24 hours at the temperature of interest. We tabulated the difference in OD_600_ between the final and initial timepoints across this 24-hour incubation for each replicate culture. The vector of these replicate measurements for each strain at a given temperature was compared to that from *S. paradoxus* with a one-tailed Wilcoxon test. In Figure S2, lines are the result of a polynomial regression on the points, created using Seaborn’s regplot in Python 3.7.

For measurements of growth at 4°C in Figure 4 and Figures S6-S8, for a given day’s worth of experiments, strains were streaked on solid plates and three to eight colonies per genotype, respectively, were pre-cultured in liquid, each as an independent biological replicate, as above. After the second back-dilution, each liquid culture was incubated for 11 days at 4°C in a rotating shaker at maximum speed in a cold room. The OD_600_ was measured on days 0, 4, 5, 7, 8, 9, 10, and 11, and also on day 6 for Figures S6-S8. To measure biomass accumulation (growth efficiency) we tabulated the difference in OD_600_ between the final and initial timepoints across this 11-day incubation for each replicate culture. Two such days’ worth of experiments were carried out for each strain in Figures S7 and S8, and one in Figures 4 and S6. We collated the measurements from all replicate culture measurements across all days for a given strain and compared each transgenic against *S. paradoxus* with a one-tailed Wilcoxon test. Separately, we fit a logistic curve to the timecourse measurements for each replicate using Scipy’s curve_fit function as a part of Scipy’s optimize package (Python 3.7). Bounds for the parameters of the logistic equation (the carrying capacity, growth rate, and time to half-maximal growth) were constrained to the range -1.0 to 10.0, and the Trust Region Reflective algorithm was used to find the best fit. We collated the growth rate estimates from all replicate culture measurements across all days for a given strain and compared each transgenic against *S. paradoxus* with a one-tailed Wilcoxon test.

### Viability assays

For the survey of viability phenotypes at high temperatures across environmental isolates in Figure S4, strains were streaked out and four colonies of each were pre-cultured in liquid as for 39°C growth above, except that the initial pre-culture to achieve saturation lasted 48 hours. Each pre-culture was back-diluted into 10 mL of YPD to reach an OD_600_ of 0.05 and then cultured for 24 hours at the temperature of interest (35°C -38°C). The OD_600_ at the end of this timecourse was measured for each such replicate culture. Then to measure viability for each, we diluted aliquots from the culture in a 1:10 series and spotted 3µL of each dilution for growth on a solid YPD plate. After incubation at 28°C for two days, we used the dilution corresponding to the most dense spot that was not a lawn for the final report of viability: we counted the number of colonies in each of the two technical replicate spots, formulated the number of colony-forming units per mL of undiluted culture (CFU/mL), and divided this ratio by the OD_600_ we had measured at the end of the liquid timecourse, to account for differences in the number of dead cells that contribute to the latter. At a given temperature, the vector of viability measurements across all replicate liquid cultures for all *S. cerevisiae* strains was compared to that for all *S. paradoxus* strains with a one-tailed Wilcoxon test.

For the comparison of viability between strains during log-phase growth at 39°C in Figure 3A, strains were streaked out and three colonies of each were pre-cultured at 28°C, back-diluted, and cultured at 39°C, as for the 39°C growth assays above. After 24 hours of incubation at 39°C, for each such replicate culture, spotting of dilutions and colony counting to yield CFU/mL/OD_600_ for each replicate liquid culture was done as above, except that we used two technical replicate spotting assay replicates for each culture, taking the average across them as the final report of viability. The vector of these viability measurements across replicates for a given strain was compared to that from *S. paradoxus* with a one-tailed Wilcoxon test.

For the comparison of viability between strains in stationary phase at 39°C in Figure 3B, strains were streaked out and three colonies of each were pre-cultured at 28°C as for the 39°C growth assays above, except that that the initial pre-culture to achieve saturation lasted 72 hours. Each such replicate culture was then incubated (without back-dilution) at 39°C for 24 hours, after which spotting, colony counting, and statistical testing were as for Figure 3A.

For the comparison of viability between strains during log-phase growth at 28°C in Figure S5A, strains were streaked out and four colonies of each were pre-cultured at 28°C and back-diluted as for the 39°C growth assays above; each back-diluted replicate culture was incubated at 28°C for 24 hours, after which spotting and colony counting was as above. For the comparison of viability between strains in stationary phase at 28°C in Figure S5B, strains were streaked out as above, and three colonies of each were inoculated into liquid YPD at 28°C and incubated for 96 hours, after which spotting, colony counting, and statistical testing were as for Figure 3A, except that a two-sided Wilcoxon tests were performed.

For the comparison of viability between environmental strains during log-phase growth at 4°C (Figure S6), aliquots from cultures set up for cold growth assays (see above) were taken at day 8 of the cold timecourse for spotting, colony counting, and statistical testing as for Figure 3A.

### Epistasis analysis

For Figure 2A, we calculated the growth phenotype at 39°C of the strain harboring all eight *S. cerevisiae* loci in the *S. paradoxus* background as expected under a model of independent locus effects as follows. We first tabulated the mean growth efficiency at 39°C of each isogenic strain in turn with just one gene swapped in from *S. cerevisiae* and the analogous mean for *S. paradoxus*, and took the difference between them, representing the mean effect of the respective swap; we then summed the latter effect values across all eight loci. Error bars were calculated by bootstrapping as follows. For each locus, we generated a random sample of the replicate measurements of the growth efficiency of the respective swap strain at 39°C with replacement, and took the mean; we calculated an analogous mean from a random sample of replicates of the *S. paradoxus* wild-type; and we took the difference between these means, representing one bootstrapped estimate of the effect of the swap. We then took the sum of such effects across all loci, representing one bootstrap’s worth of the estimate of the eight-gene transgenic’s phenotype under the additive model. We repeated this procedure 10,000 times to set up a distribution of the estimated sum, and we identified the values corresponding to the 68% confidence interval.

For a given panel of Figure 2B, the left-hand point on the plot reports the mean effect of the *S. cerevisiae* allele of the respective gene when swapped alone into *S. paradoxus*. For this we tabulated the mean growth efficiency at 39°C of this single-gene transgenic across all replicates and the analogous mean across all replicates of *S. paradoxus*, and took the difference between them. For the error bar, we generated a random sample of the replicate measurements of the growth efficiency of the respective swap strain at 39°C with replacement, and took the mean; we calculated an analogous mean from a random sample of replicates of the *S. paradoxus* wild-type; and we took the difference between these means, representing one bootstrapped estimate of the effect of the swap. We repeated this procedure 10,000 times to set up a distribution of the effect value, and we identified the values corresponding to the 68% confidence interval. The right-hand point on the plot reports the mean effect of the *S. cerevisiae* allele of the respective gene when swapped into a multi-genic strain of the *S. paradoxus* background in the series culminating in the eight-fold transgenic, in the order of Figure 1; call the strain before and after the replacement of the gene of interest *X-1* and *X*, respectively. We tabulated the mean growth efficiency at 39°C across all replicates of strain *X* and the analogous quantity for strain *X-1*, and took the difference between them. Error bars were calculated by bootstrapping as above.

## Supplementary table captions

**Table S1. Statistical analyses of growth at 39°C and 28°C**. (A) Each cell reports the results of a one-sided Wilcoxon test comparing growth efficiency at 39°C between the indicated strains, in Figure 1 of the main text. Multiple testing was corrected for using the Benjamini-Hochberg method. (B) Data are as in A except that analysis was of growth efficiency at 28°C from Figure S1 and two-sided Wilcoxon tests were applied.

**Table S2. Parameters used in regressions**. Each row reports fitted values of the indicated parameters from the polynomial regression (Figure S2) or logistic regression (all other figures) of growth measurements of the indicated strain for the indicated figure. For the logistic regression, K, R, and x_0_ are the carrying capacity, logistic growth rate, and the sigmoidal midpoint, respectively.

**Table S3. Strains used in this study. A**. Wild-type diploid strains, including those used as parents of allele-replacement transgenesis; SGRP, the Saccharomyces Genome Resequencing Project, version 2. **B**. Allele replacement strains in *S. paradoxus* Z1 diploid homozygote backgrounds. In genotype notes, *e*.*g*., in an *S. paradoxus* background, ΔYFG(-X to +Y)::scYFG(-Z to +W) indicates that in *S. paradoxus* Z1, bases -X to +Y from gene YFG have been removed and replaced by bases -Z to +W of the allele of YFG from the indicated *S. cerevisiae* strain. Positive coordinates count in the 5’ to 3’ direction from the start codon (+1 corresponds to the A in the ATG), and negative coordinates count in the 3’ to 5’ direction from the start codon (−1 corresponds to the base directly 5’ of the ATG). In cases where the replacement extended into a region of 100% conservation between species, the position of the last divergent nucleotide is shown.

## Supplementary figure captions

**Figure S1. Allelic variation at thermotolerance loci has little growth impact at 28°C**. Data and symbols are as in Figure 1 of the main text except that growth was measured at 28°C. Statistical analyses are reported in Table S1B.

**Figure S2. S. cerevisiae alleles of thermotolerance loci jointly improve growth at temperatures from 37°C -39°C**. In a given panel, each trace reports growth efficiency, the cell density after a 24-hour incubation at the indicated temperature as a difference from the starting density, of the wild-type of the indicated species or the S. paradoxus strain harboring eight thermotolerance loci from S. cerevisiae (8x swap). Lines are the result of a polynomial regression on the points (Table S2). *, Wilcoxon p ≤ 0.0404.

**Figure S3. *S. cerevisiae* alleles of thermotolerance loci individually improve growth at high temperature**. Data and symbols are as in the main panel of Figure 1 except that each column reports results from the indicated wild-type strain or a strain of *S. paradoxus* harboring the *S. cerevisiae* allele of the indicated single gene and the *y*-axis is log-scaled. All comparisons to *S. paradoxus* had one-sided Wilcoxon *p* < 0.01 after correction for multiple testing.

**Figure S4. Environmental isolates of Saccharomyces spp. differ in heat survival**. Each cell reports viability after heat treatment of the indicated strain and species. Each cell reports the number of colonies formed on solid medium from 1 mL of heat-treated liquid culture in logarithmic growth, normalized by the turbidity of the latter. *S. paradoxus* Z1, N17, and A12 are isolates from UK, Russia, and Quebec, respectively; *S. cerevisiae* strains DBVPG1373, DBVPG1788, YPS128 are from the Netherlands, Finland, and Pennsylvania, respectively. Viability was different between species at Wilcoxon *p* < 0.00002 for all temperatures.

**Figure S5. Allelic variation at thermotolerance loci has no impact on viability at 28°C**. Data and symbols are as in Figure 3 of the main text except that liquid incubations were at 28°C. In no case was the respective measurement for a given strain significantly different from the analogous quantity for *S. paradoxus* at two-sided Wilcoxon *p* < 0.05.

**Figure S6. Environmental isolates of *Saccharomyces spp*. differ in ability to grow at 4°C**. (A) Each trace reports a timecourse of growth at 4°C of the wild-type of the indicated strain. For a given strain, points on a given day report biological replicates; lines report the average fit from a logistic regression across replicates (Table S2). YPS128, a North American *S. cerevisiae* known to have recently acquired freeze-thaw resistance as a derived character distinct from the ancestral program (Will et al., 2010), is shown in faint blue. (B) The *y*-axis reports growth rate, in units of cell density (optical density, OD) per day, from the average logistic fit of the timecourse in (A) for the indicated strain. (C) The *y*-axis reports, for day 10 of the timecourse in (A) for the indicated strain, growth efficiency, the cell density after a 10-day incubation at 4°C as a difference from the starting density. In (B) and (C), Black points report individual biological replicates. White dots report means. Boxes span the interquartile range; whiskers are 1.5 times the interquartile range, and do not report outliers. Comparisons of growth rate and efficiency at 4°C between *S. paradoxus* strains and *S. cerevisiae* strains yielded Wilcoxon *p* ≤ 0.00003 and *p* ≤ 0.002, respectively.

**Figure S7. Growth effects at 4°C of *S. cerevisiae* alleles of individual thermotolerance loci**. (A) Data and symbols are as in Figure S3 except that the *y*-axis reports growth efficiency after a 10-day incubation at 4°C. (B) Data are as in (A) except that the y-axis reports growth rate, in units of cell density (optical density, OD) per day, from the average logistic fit of the timecourse for the indicated strain (Table S2). * and **, one-sided Wilcoxon *p* ≤ 0.05 and 0.01, respectively.

**Figure S8. Joint growth effects at 4°C of S. cerevisiae alleles of subsets of thermotolerance loci**. (A) Data and symbols are as in the main panel of Figure 1 except that the y-axis reports growth efficiency after a 10-day incubation at 4°C. (B) Data and symbols are as in (A) except that the y-axis reports growth rate, in units of cell density (optical density, OD) per day, from the average logistic fit of the timecourse for the indicated strain (Table S2). * and **, one-sided Wilcoxon p ≤ 0.05 and 0.01 respectively.

**Figure S9. Allelic variation at thermotolerance loci has little impact on viability at 4°C**. Data and symbols are as in Figure 3A of the main text, except that liquid incubations were at 4°C, and measurements were taken at day 8 of the growth timecourse. *, One-sided Wilcoxon *p* ≤ 0.05.

