## Supplementary material for "Joint effects of genes underlying a temperature specialization tradeoff in yeast": Tables S1-S3; Figures S1-S9

**A**

| | 1x swap | 2x swap | 3x swap | 4x swap | 5x swap | 6x swap | 7x swap | 8x swap | *S. cerevisiae* |
|---|---|---|---|---|---|---|---|---|---|
| ***S. paradoxus*** | 0.000005 | 0.000006 | 0.000010 | 0.000000 | 0.000000 | 0.000000 | 0.000000 | 0.000000 | 0.000000 |
| **1x swap** | | 0.088928 | 0.517759 | 0.007992 | 0.000003 | 0.000001 | 0.000000 | 0.000000 | 0.000000 |
| **2x swap** | | | 0.565374 | 0.110054 | 0.000443 | 0.000009 | 0.000000 | 0.000000 | 0.000000 |
| **3x swap** | | | | 0.077462 | 0.001568 | 0.000048 | 0.000000 | 0.000000 | 0.000000 |
| **4x swap** | | | | | 0.005735 | 0.000112 | 0.000000 | 0.000000 | 0.000000 |
| **5x swap** | | | | | | 0.037984 | 0.000010 | 0.000000 | 0.000000 |
| **6x swap** | | | | | | | 0.014711 | 0.001019 | 0.000000 |
| **7x swap** | | | | | | | | 0.037984 | 0.000000 |
| **8x swap** | | | | | | | | | 0.000000 |

**B**

| | 1x swap | 2x swap | 3x swap | 4x swap | 5x swap | 6x swap | 7x swap | 8x swap | *S. cerevisiae* |
|---|---|---|---|---|---|---|---|---|---|
| ***S. paradoxus*** | 0.860008 | 0.860008 | 0.860008 | 0.860008 | 0.860008 | 0.860008 | 0.860008 | 0.960778 | 0.860008 |
| **1x swap** | | 1.000000 | 0.860008 | 0.860008 | 1.000000 | 0.960778 | 0.860008 | 0.860008 | 0.860008 |
| **2x swap** | | | 0.860008 | 0.860008 | 1.000000 | 1.000000 | 0.960778 | 0.860008 | 0.860008 |
| **3x swap** | | | | 0.860008 | 0.860008 | 1.000000 | 0.860008 | 0.860008 | 0.860008 |
| **4x swap** | | | | | 0.860008 | 0.860008 | 0.860008 | 1.000000 | 0.860008 |
| **5x swap** | | | | | | 1.000000 | 0.960778 | 0.860008 | 0.860008 |
| **6x swap** | | | | | | | 0.860008 | 0.860008 | 0.860008 |
| **7x swap** | | | | | | | | 1.000000 | 0.860008 |
| **8x swap** | | | | | | | | | 0.860008 |

**Table S1. Statistical analyses of growth at 39°C and 28°C.** (A) Each cell reports the results of a one-sided Wilcoxon test comparing growth efficiency at 39°C between the indicated strains, in Figure 1 of the main text. Multiple testing was corrected for using the Benjamini-Hochberg method. (B) Data are as in A except that analysis was of growth efficiency at 28°C from Figure S1 and two-sided Wilcoxon tests were applied.

**Equation:**

$$f(x) = \frac{K}{1 + e^{-R(x-x_0)}}$$

| K | R | $x_0$ | Strain | Figure |
|---|---|---|---|---|
| 3.837976 | 0.683081 | 7.611606 | *S. paradoxus* Z1 | 4 |
| 3.76661 | 0.64131 | 7.97884 | 8x swap | 4 |
| 3.86467 | 0.635313 | 7.994714 | S. *cerevisiae* DBVPG1737 | 4 |
| 4.459902 | 0.718399 | 6.68077 | *S. paradoxus* Z1 | S6 |
| 5.047729 | 0.680221 | 6.927175 | *S. paradoxus* N17 | S6 |
| 4.592433 | 0.652344 | 7.044108 | *S. paradoxus* IFO1804 | S6 |
| 3.297667 | 0.727683 | 6.213845 | *S. paradoxus* DBVPG6304 | S6 |
| 4.645203 | 0.673755 | 6.944962 | *S. paradoxus* A12 | S6 |
| 3.679257 | 0.375646 | 10 | S. *cerevisiae* DBVPG1737 | S6 |
| 2.258294 | 0.286145 | 10 | S. *cerevisiae* YS2 | S6 |
| 2.841081 | 0.430244 | 7.594446 | S. *cerevisiae* DBVPG1788 | S6 |
| 2.272461 | 0.420579 | 10 | S. *cerevisiae* UWOPS03.461.4 | S6 |

| | | | | |
|---|---|---|---|---|
| 5.191947 | 0.527724 | 8.510967 | S. *cerevisiae* YPS128 | S6 |
| 4.494999 | 0.79778 | 6.058288 | *S. paradoxus* Z1 | S7 |
| 4.205824 | 0.756281 | 6.034304 | AFG2 1x swap | S7 |
| 4.403604 | 0.818116 | 6.105421 | APC1 1x swap | S7 |
| 4.244261 | 0.793644 | 6.076612 | CEP3 1x swap | S7 |
| 3.83629 | 0.782956 | 5.960992 | DYN1 1x swap | S7 |
| 4.45455 | 0.824422 | 5.984797 | ESP1 1x swap | S7 |
| 4.134131 | 0.820255 | 5.811504 | MYO1 1x swap | S7 |
| 4.15255 | 0.802674 | 5.997295 | SCC2 1x swap | S7 |
| 4.258058 | 0.753425 | 5.976158 | TAF2 1x swap | S7 |
| 4.64392 | 0.410473 | 10 | S. *cerevisiae* DBVPG1737 | S7 |
| 4.201285 | 0.738212 | 6.533661 | *S. paradoxus* Z1 | S8 |
| 3.742483 | 0.689709 | 6.53867 | 1x swap | S8 |
| 3.56338 | 0.657921 | 6.616394 | 2x swap | S8 |
| 3.504207 | 0.699821 | 6.478092 | 3x swap | S8 |
| 3.43079 | 0.680154 | 6.430598 | 4x swap | S8 |
| 3.588065 | 0.676851 | 6.46178 | 5x swap | S8 |
| 3.485715 | 0.7081 | 6.394901 | 6x swap | S8 |

| | | | | |
|---|---|---|---|---|
| 3.511028 | 0.686568 | 6.490653 | 7x swap | S8 |
| 3.650739 | 0.637369 | 6.688769 | 8x swap | S8 |
| 3.583343 | 0.421157 | 10 | S. *cerevisiae* DBVPG1737 | S8 |
| **Equation:** $f(x) = ax^2 + bx + c$ | | | | |
| **a** | **b** | **c** | **Strain** | **Figure** |
| 7.76E-01 | -6.05E+01 | 1.18E+03 | *S. paradoxus* Z1 | S2 |
| 7.80E-01 | -6.08E+01 | 1.18E+03 | 8x swap | S2 |
| 9.38E-01 | -7.24E+01 | 1.40E+03 | *S. cerevisiae* DBVPG1737 | S2 |

**Table S2. Parameters used in regressions.** Each row reports fitted values of the indicated parameters from the polynomial regression (Figure S2) or logistic regression (all other figures) of growth measurements of the indicated strain for the indicated figure. For the logistic regression, K, R, and $x_0$ are the carrying capacity, logistic growth rate, and the sigmoidal midpoint, respectively.

| Species | Strain Name | Background | Genotype/Details (Population designation from Liti et al., 2009; Peter et al., 2018) | Source | Allele donor strain |
| --- | --- | --- | --- | --- | --- |
| **A.** | | | | | |
| S. par | Z1 | | Tree bark isolate from Silwood Park, UK (European) | SGRP | |
| S. par | N17 | | Oak exudate isolate from Russia (European) | SGRP | |
| S. par | A12 | | Oak isolate from Quebec, CA (American) | SGRP | |
| S. par | IFO1804 | | Oak bark isolate from Japan (Far Eastern) | SGRP | |
| S. par | DBVPG6304 | | *Drosophila* from Yosemite, USA (American) | SGRP | |
| S. cer | DBVPG1737 | | Soil isolate from Netherlands (Wine/European) | SGRP | |
| S. cer | DBVPG1788 | | Soil isolate from Finland (Wine/European) | SGRP | |
| S. cer | YPS128 | | Woodland isolate from Pennsylvania, USA (North American) | SGRP | |

| | | | | |
|---|---|---|---|---|
| S. cer | UWOPS03.461.4 | | Bertam palm nectar isolate from Malaysia (Mosaic) | SGRP |
| S. cer | YS2 | | Baker's strain from Australia (Baking) | SGRP |

**B.**

| | | | | | |
|---|---|---|---|---|---|
| S. par | CW223 | Z1 | Δ*DYN1*(-231 to 12377):: sc*DYN1*(-243 to 12377) | Weiss et al. 2018 | DBVPG1373 |
| S. par | CW382 | CW223 | Δ*DYN1*(-231 to 12377):: sc*DYN1*(-243 to 12377)<br><br>Δ*MYO1*(-907 to 5826):: sc*MYO1*(-903 to 5824) | This study | DBVPG1373 |
| S. par | CW415 | CW382 | Δ*DYN1*(-231 to 12377):: sc*DYN1*(-243 to 12377)<br><br>Δ*MYO1*(-907 to 5826):: sc*MYO1*(-903 to 5824)<br><br>Δ*SCC2*(-456 to 4556):: sc*SCC2*(-441 to 4556) | This study | DBVPG1373 |
| S. par | CW417 | CW415 | Δ*DYN1*(-231 to 12377):: sc*DYN1*(-243 to 12377)<br><br>Δ*MYO1*(-907 to 5826):: sc*MYO1*(-903 to 5824)<br><br>Δ*SCC2*(-456 to 4556):: sc*SCC2*(-441 to 4556) | This study | DBVPG1373 |

| | | | | | |
|---|---|---|---|---|---|
| | | | Δ*APC1*(-547 to 5280)::<br>sc*APC1*(-554 to 5280) | | |
| S. par | FZ3 | CW417 | Δ*DYN1*(-231 to 12377)::<br>sc*DYN1*(-243 to 12377)<br>Δ*MYO1*(-907 to 5826)::<br>sc*MYO1*(-903 to 5824)<br>Δ*SCC2*(-456 to 4556)::<br>sc*SCC2*(-441 to 4556)<br>Δ*APC1*(-547 to 5280)::<br>sc*APC1*(-554 to 5280)<br>Δ*AFG2*(-293 to 2391)::<br>sc*AFG2*(-270 to 2391) | This study | DBVPG1373 |
| S. par | FZ4 | FZ3 | Δ*DYN1*(-231 to 12377)::<br>sc*DYN1*(-243 to 12377)<br>Δ*MYO1*(-907 to 5826)::<br>sc*MYO1*(-903 to 5824)<br>Δ*SCC2*(-456 to 4556)::<br>sc*SCC2*(-441 to 4556)<br>Δ*APC1*(-547 to 5280)::<br>sc*APC1*(-554 to 5280)<br>Δ*AFG2*(-293 to 2391)::<br>sc*AFG2*(-270 to 2391) | This study | DBVPG1373 |

| | | | | | |
|---|---|---|---|---|---|
| | | | ΔESP1(-418 to 4962)::<br>scESP1(-439 to 4963) | | |
| S. par | FZ15 | FZ4 | Δ*DYN1*(-231 to 12377)::<br>sc*DYN1*(-243 to 12377)<br>Δ*MYO1*(-907 to 5826)::<br>sc*MYO1*(-903 to 5824)<br>Δ*SCC2*(-456 to 4556)::<br>sc*SCC2*(-441 to 4556)<br>Δ*APC1*(-547 to 5280)::<br>sc*APC1*(-554 to 5280)<br>Δ*AFG2*(-293 to 2391)::<br>sc*AFG2*(-270 to 2391)<br>ΔESP1(-418 to 4962)::<br>scESP1(-439 to 4963)<br>Δ*CEP3*(-187 to 1816)::<br>sc*CEP3*(-194 to 1816) | This study | DBVPG1373 |
| S. par | FZ22 | FZ15 | Δ*DYN1*(-231 to 12377)::<br>sc*DYN1*(-243 to 12377)<br>Δ*MYO1*(-907 to 5826)::<br>sc*MYO1*(-903 to 5824)<br>Δ*SCC2*(-456 to 4556)::<br>sc*SCC2*(-441 to 4556)<br>Δ*APC1*(-547 to 5280)::<br>sc*APC1*(-554 to 5280) | This study | DBVPG1373 |

| | | | |
|---|---|---|---|
| | | | Δ*AFG2*(-293 to 2391)::<br>sc*AFG2*(-270 to 2391)<br><br>ΔESP1(-418 to 4962)::<br>scESP1(-439 to 4963)<br><br>Δ*CEP3*(-187 to 1816)::<br>sc*CEP3*(-194 to 1816)<br><br>Δ*TAF2*(-806 to 4348)::<br>sc*TAF2*(-774 to 4357) |

**Table S3. Strains used in this study. A.** Wild-type diploid strains, including those used as parents of allele-replacement transgenesis; SGRP, the Saccharomyces Genome Resequencing Project, version 2. **B.** Allele replacement strains in *S. paradoxus* Z1 diploid homozygote backgrounds. In genotype notes, *e.g.*, in an *S. paradoxus* background, ΔYFG(-X to +Y)::scYFG(-Z to +W) indicates that in *S. paradoxus* Z1, bases -X to +Y from gene YFG have been removed and replaced by bases -Z to +W of the allele of YFG from the indicated *S. cerevisiae* strain. Positive coordinates count in the 5' to 3' direction from the start codon (+1 corresponds to the A in the ATG), and negative coordinates count in the 3' to 5' direction from the start codon (-1 corresponds to the base directly 5' of the ATG). In cases where the replacement extended into a region of 100% conservation between species, the position of the last divergent nucleotide is shown.

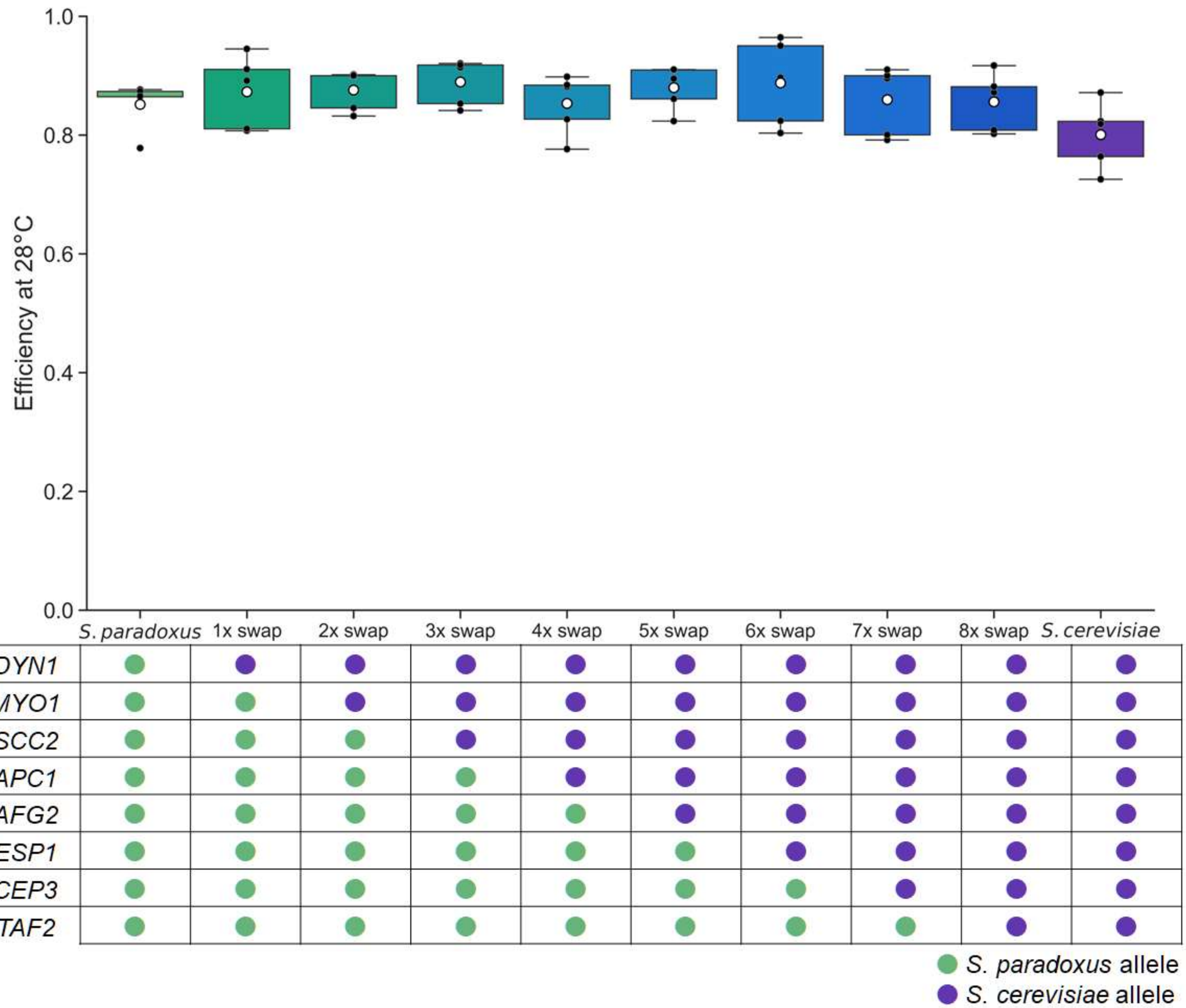

**Figure S1. Allelic variation at thermotolerance loci has little growth impact at 28°C.** Data and symbols are as in Figure 1 of the main text except that growth was measured at 28°C. Statistical analyses are reported in Table S1B.

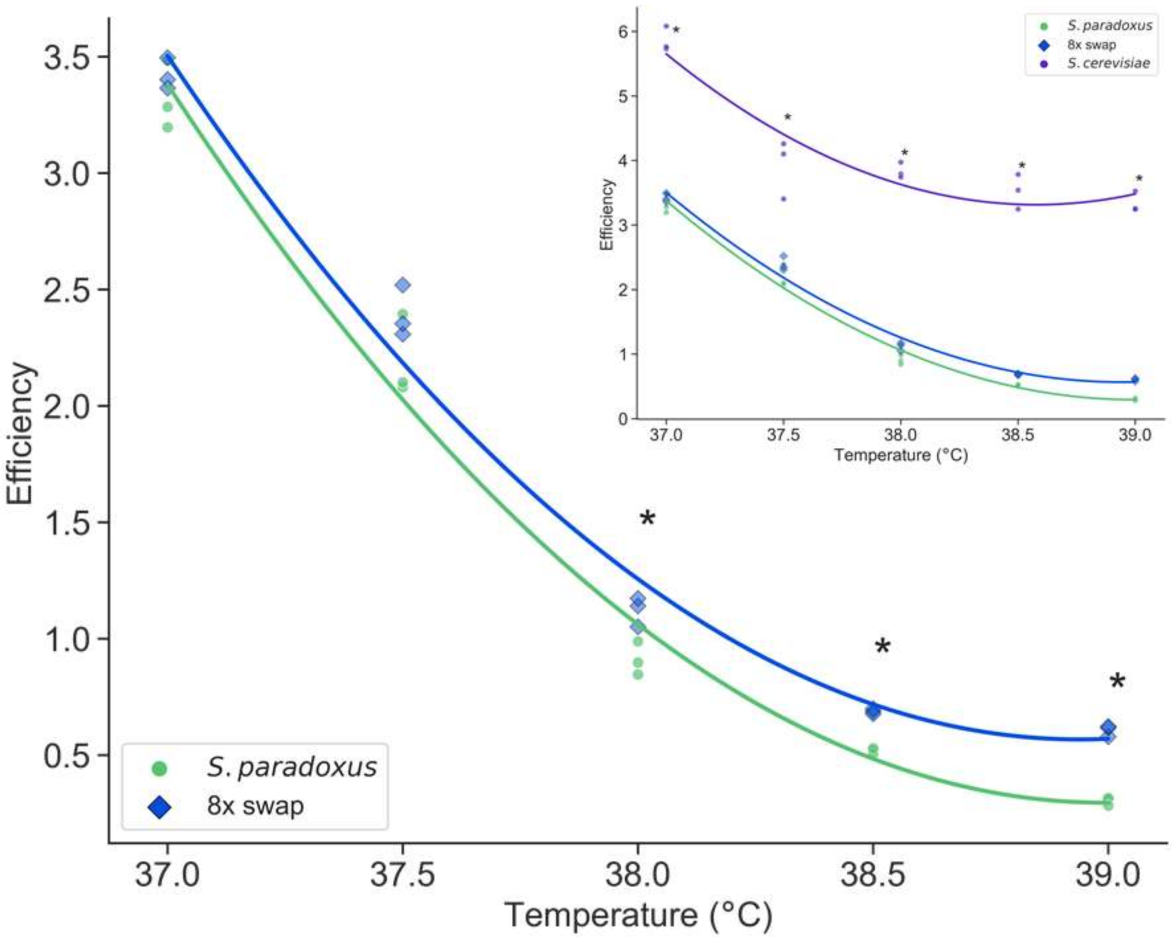

**Figure S2. *S. cerevisiae* alleles of thermotolerance loci jointly improve growth at temperatures from 37°C - 39°C.** In a given panel, each trace reports growth efficiency, the cell density after a 24-hour incubation at the indicated temperature as a difference from the starting density, of the wild-type of the indicated species or the *S. paradoxus* strain harboring eight thermotolerance loci from *S. cerevisiae* (8x swap). Lines are the result of a polynomial regression on the points (Table S2). *, Wilcoxon $p \leq 0.0404$.

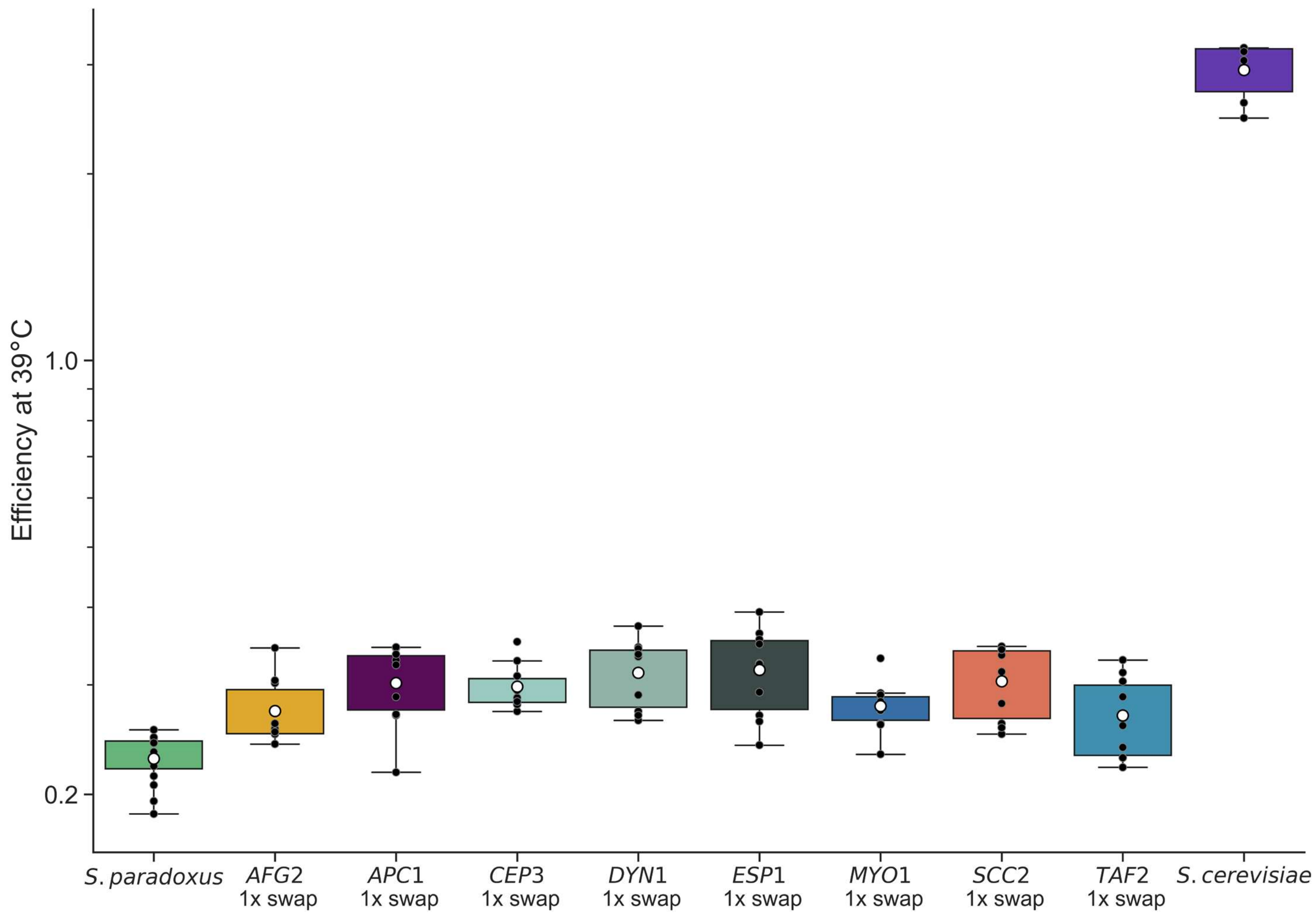

**Figure S3. *S. cerevisiae* alleles of thermotolerance loci individually improve growth at high temperature.** Data and symbols are as in the main panel of Figure 1 except that each column reports results from the indicated wild-type strain or a strain of *S. paradoxus* harboring the *S. cerevisiae* allele of the indicated single gene and the *y*-axis is log-scaled. All comparisons to *S. paradoxus* had one-sided Wilcoxon $p < 0.01$ after correction for multiple testing.

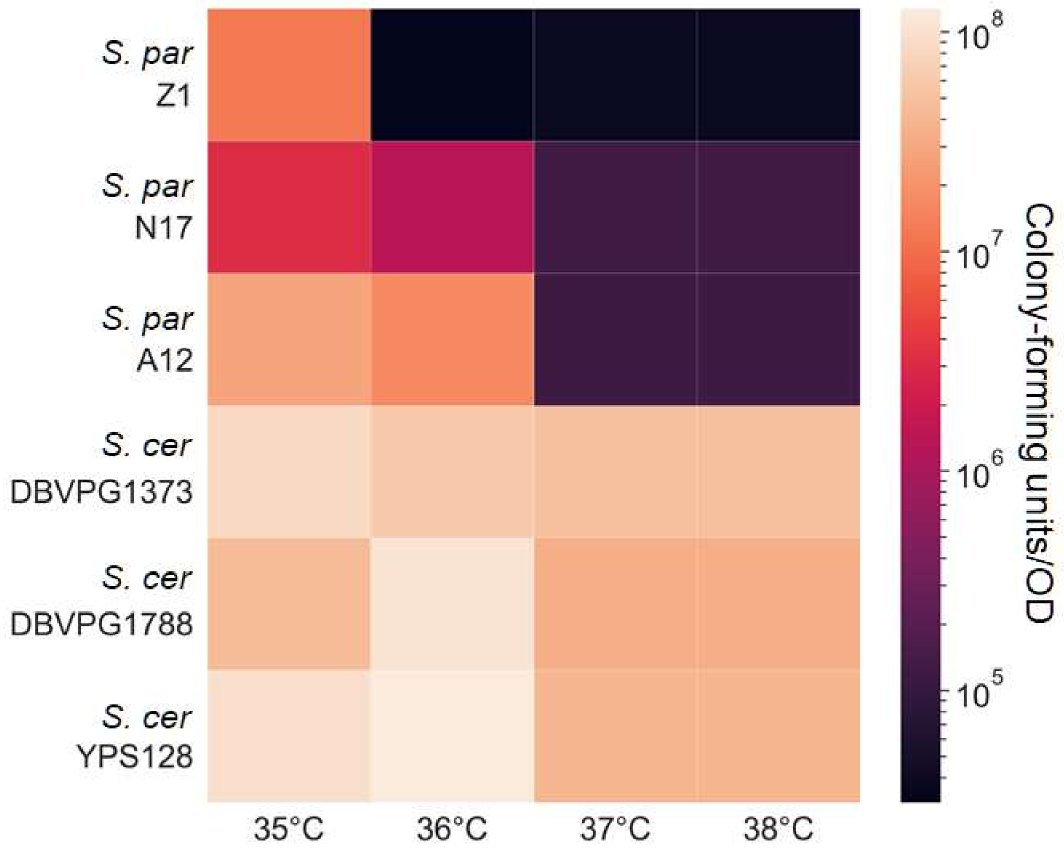

**Figure S4. Environmental isolates of Saccharomyces spp. differ in heat survival.** Each cell reports viability after heat treatment of the indicated strain and species. Each cell reports the number of colonies formed on solid medium from 1 mL of heat-treated liquid culture in logarithmic growth, normalized by the turbidity of the latter. *S. paradoxus* Z1, N17, and A12 are isolates from UK, Russia, and Quebec, respectively; *S. cerevisiae* strains DBVPG1373, DBVPG1788, YPS128 are from the Netherlands, Finland, and Pennsylvania, respectively. Viability was different between species at Wilcoxon $p < 0.00002$ for all temperatures.

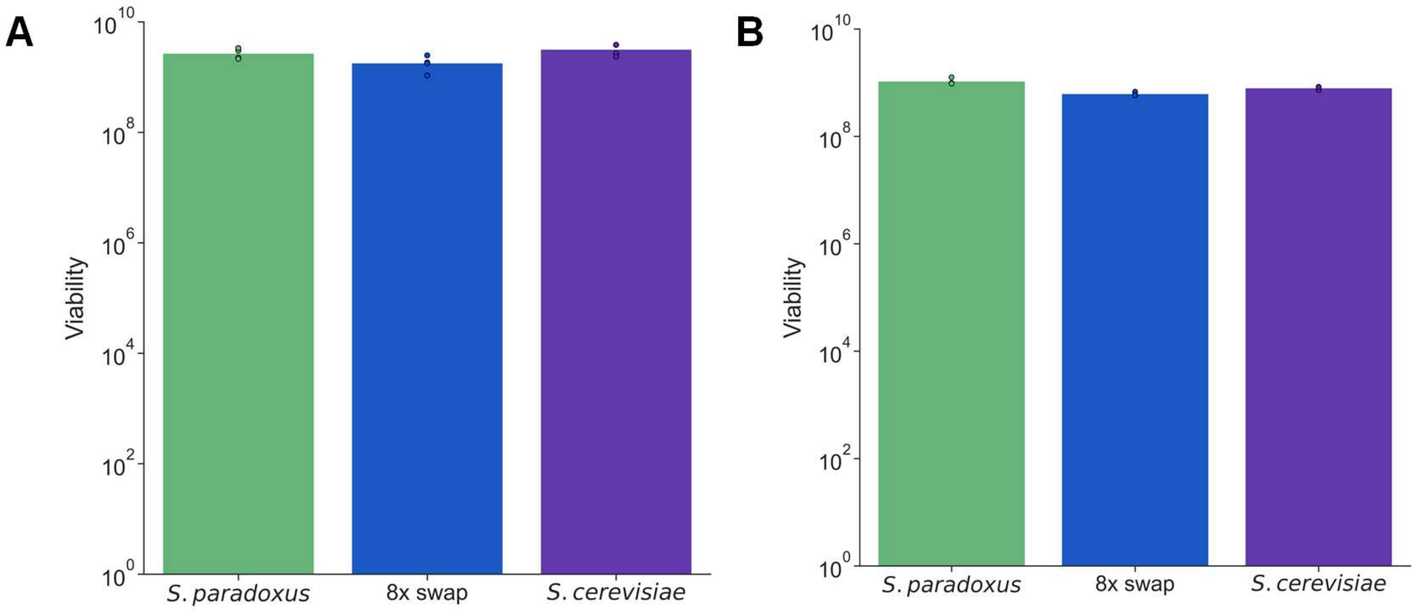

**Figure S5. Allelic variation at thermotolerance loci has no impact on viability at 28°C.** Data and symbols are as in Figure 3 of the main text except that liquid incubations were at 28°C. In no case was the respective measurement for a given strain significantly different from the analogous quantity for *S. paradoxus* at two-sided Wilcoxon $p < 0.05$.

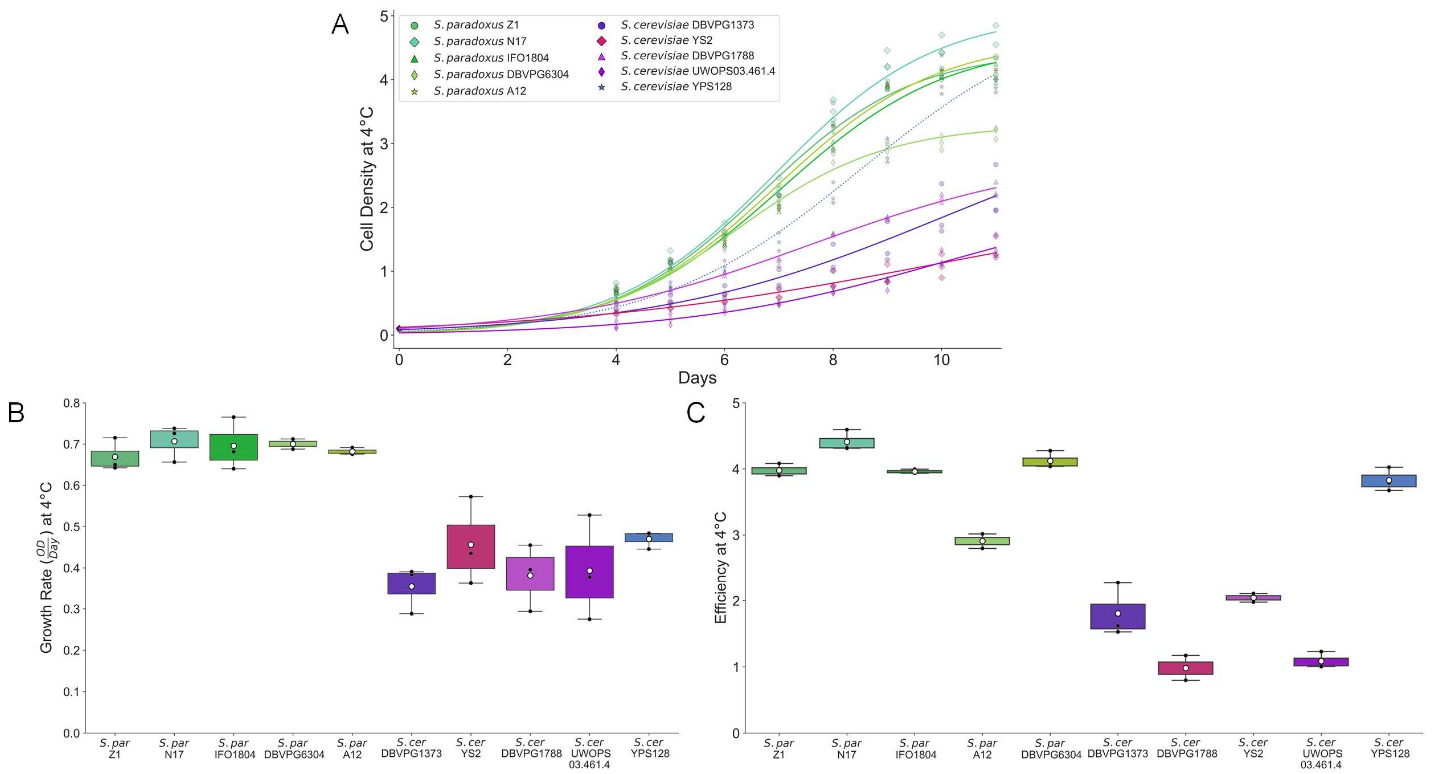

**Figure S6. Environmental isolates of *Saccharomyces spp.* differ in ability to grow at 4°C.** (A) Each trace reports a timecourse of growth at 4°C of the wild-type of the indicated strain. For a given strain, points on a given day report biological replicates; lines report the average fit from a logistic regression across replicates (Table S2). YPS128, a North American *S. cerevisiae* known to have recently acquired freeze-thaw resistance as a derived character distinct from the ancestral program (Will et al., 2010), is shown in faint blue. (B) The *y*-axis reports growth rate, in units of cell density (optical density, OD) per day, from the average logistic fit of the timecourse in (A) for the indicated strain. (C) The *y*-axis reports, for day 10 of the timecourse in (A) for the indicated strain, growth efficiency, the cell density after a 10-day incubation at 4°C as a difference from the starting density. In (B) and (C), Black points report individual biological replicates. White dots report means. Boxes span the interquartile range; whiskers are 1.5 times the interquartile range, and do not report outliers. Comparisons of growth rate and efficiency at 4°C between *S. paradoxus* strains and *S. cerevisiae* strains yielded Wilcoxon $p \leq 0.00003$ and $p \leq 0.002$, respectively.

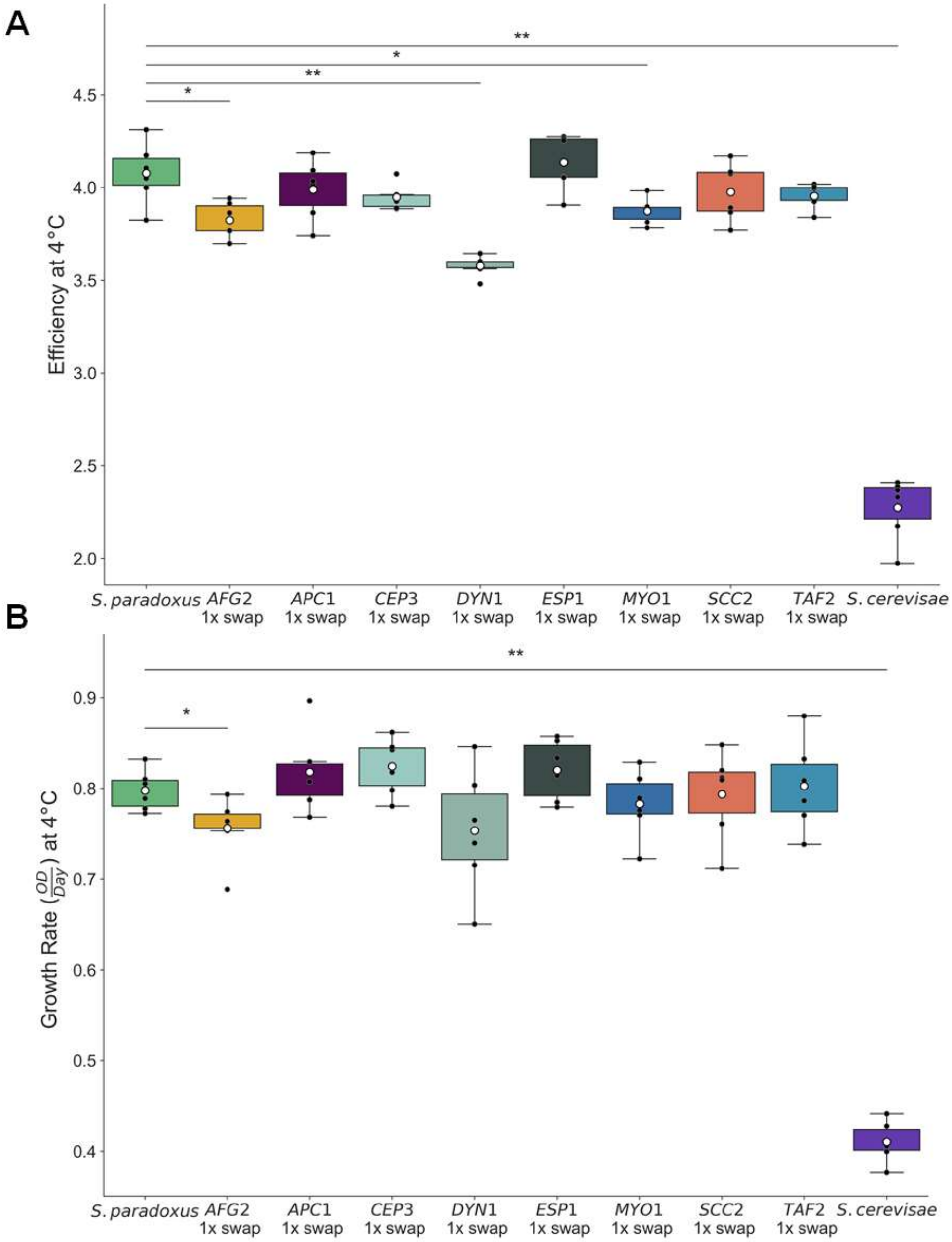

**Figure S7. Growth effects at 4°C of *S. cerevisiae* alleles of individual thermotolerance loci.** (A) Data and symbols are as in Figure S3 except that the *y*-axis reports growth efficiency after a 10-day incubation at 4°C. (B) Data are as in (A) except that the y-axis reports growth rate, in units of cell density (optical density, OD) per day, from the average logistic fit of the timecourse for the indicated strain (Table S2). * and **, one-sided Wilcoxon $p \leq 0.05$ and 0.01, respectively.

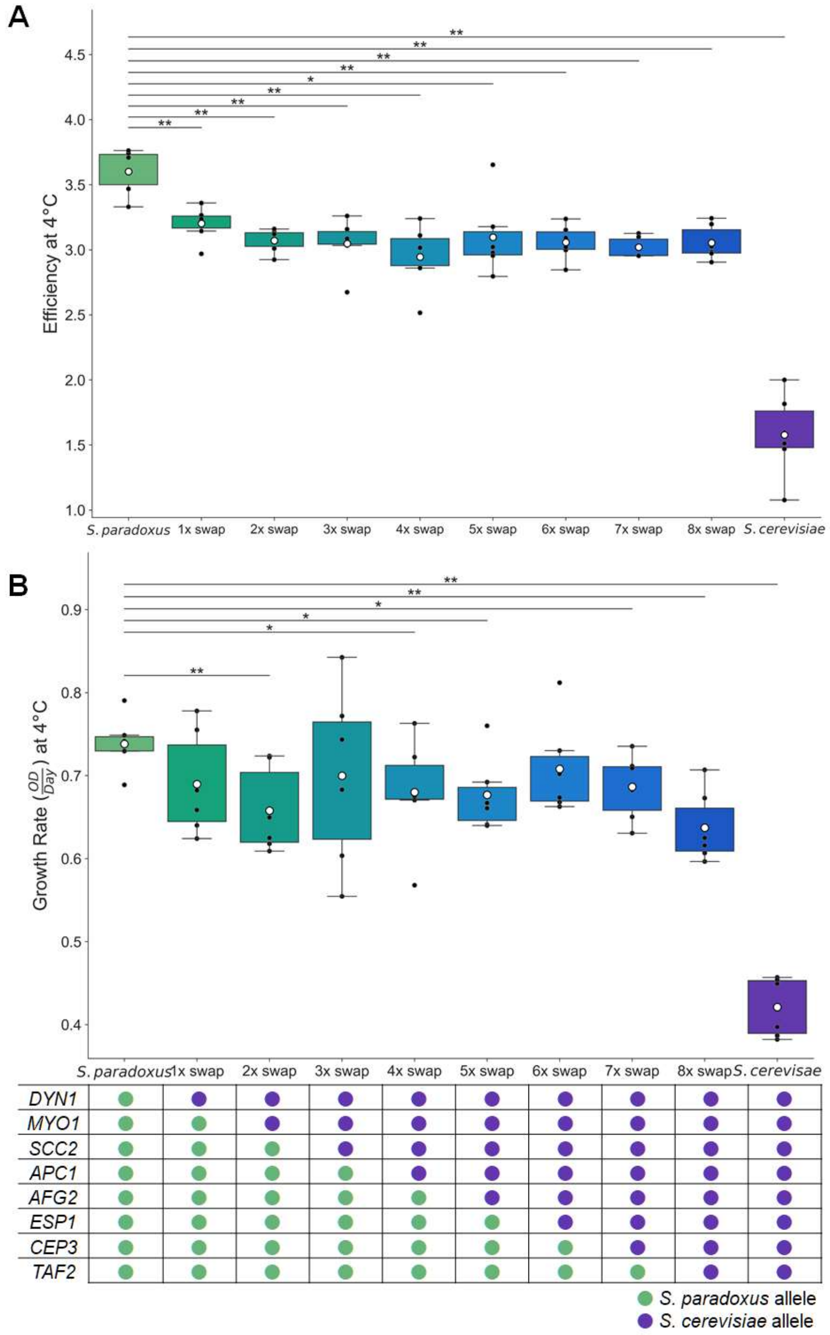

**Figure S8. Joint growth effects at 4°C of *S. cerevisiae* alleles of subsets of thermotolerance loci.** (A) Data and symbols are as in the main panel of Figure 1 except that the *y*-axis reports growth efficiency after a 10-day incubation at 4°C. (B) Data and symbols are as in (A) except that the *y*-axis reports growth rate, in units of cell density (optical density, OD) per day, from the average logistic fit of the timecourse for the indicated strain (Table S2). * and **, one-sided Wilcoxon $p \leq 0.05$ and 0.01 respectively.

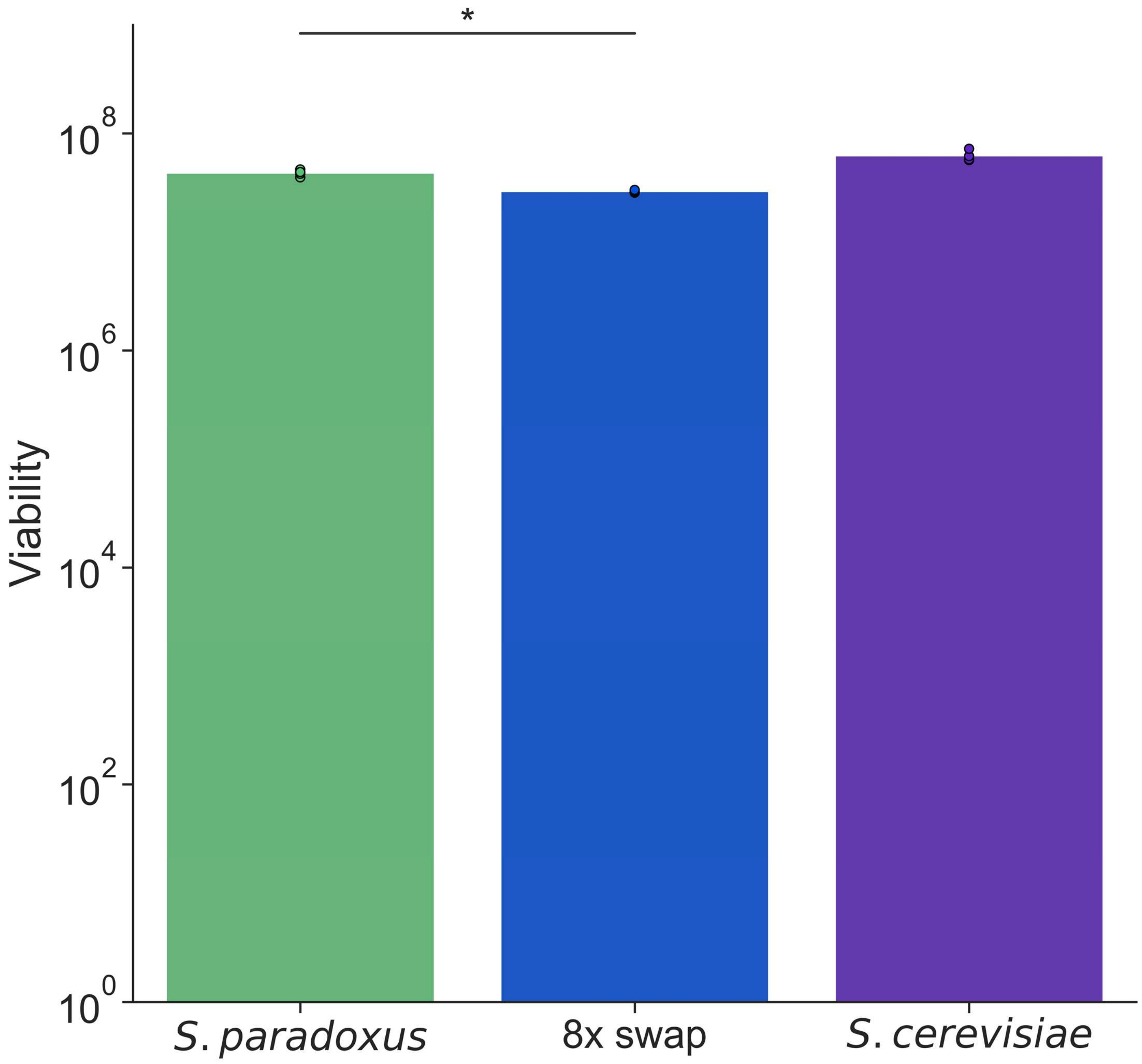

**Figure S9. Allelic variation at thermotolerance loci has little impact on viability at 4°C.** Data and symbols are as in Figure 3A of the main text, except that liquid incubations were at 4°C, and measurements were taken at day 8 of the growth timecourse. *, One-sided Wilcoxon $p \leq 0.05$.
